# Cryo-EM structure of the volume-regulated anion channel LRRC8

**DOI:** 10.1101/331207

**Authors:** Go Kasuya, Takanori Nakane, Takeshi Yokoyama, Yanyan Jia, Masato Inoue, Kengo Watanabe, Ryoki Nakamura, Tomohiro Nishizawa, Tsukasa Kusakizako, Akihisa Tsutsumi, Haruaki Yanagisawa, Naoshi Dohmae, Motoyuki Hattori, Hidenori Ichijo, Zhiqiang Yan, Masahide Kikkawa, Mikako Shirouzu, Ryuichiro Ishitani, Osamu Nureki

## Abstract

Maintenance of cell volume against osmotic change is crucial for proper cell functions, such as cell proliferation and migration. The leucine-rich repeat-containing 8 (LRRC8) proteins are anion selective channels, and were recently identified as pore components of the volume-regulated anion channels (VRACs), which extrude anions to decrease the cell volume upon cell-swelling. Here, we present the human LRRC8A structure, determined by a single-particle cryo-electron microscopy analysis. The sea anemone-like structure represents a trimer of dimers assembly, rather than a symmetrical hexameric assembly. The four-spanning transmembrane region has a gap junction channel-like membrane topology, while the LRR region containing 15 leucine-rich repeats forms a long twisted arc. The channel pore is along the central axis and constricted on the extracellular side, where the highly conserved polar and charged residues at the tip of the extracellular helix contribute to the anion and other osmolyte permeability. Comparing the two structural populations facilitated the identification of both compact and relaxed conformations, suggesting that the LRR region is flexible and mobile with rigid-body motions, which might be implicated in structural transitions upon pore opening. Overall, our structure provides a framework for understanding the molecular mechanisms of this unique class of ion channels.

## Introduction

Cell volume regulation in response to changes in the extracellular osmotic pressure is a fundamental homeostatic mechanism for all organisms^1–3^. Alteration of the cell volume leads to numerous physiological processes, such as cell proliferation and migration. A continuous decrease in the cell volume under isotonic conditions is related to the activation of the programmed cell death pathway, and is known as apoptotic volume decrease (AVD)^4^. Various proteins including membrane ion channels and transporters, as well as cytoplasmic kinases, are essential components for cell volume regulation. Among these proteins, the volume regulated anion channel (VRAC) is a particularly interesting type of ion channel^5^. VRAC is activated by cell-swelling and releases Cl? ions or other osmolytes on the extracellular side and concurrently ejects water molecules, leading to a decrease in the cell volume. This process, called regulatory volume decrease (RVD), is observed in almost all types of eukaryotic cells.

The molecular entity of VRAC was unknown until quite recently, although extensive experiments have been performed over the past decades, and several characteristics of VRAC, including substrate selectivity, activation upon cell-swelling, outward rectification, and inactivation kinetics, were functionally characterized^6,7^. In 2014, by combining high-throughput RNAi screening with electrophysiological recordings, two independent groups successfully identified the leucine-rich repeat-containing 8A (LRRC8A) protein, which when mutated causes an innate immune disease, agammaglobulinemia^8^, as the essential component of VRAC^9,10^. The LRRC8A protein belongs to the LRRC8 gene family, which includes five members (LRRC8A-E) with high amino-acid sequence similarity. All of the LRRC8 isoforms consist of about 800-850 amino acid residues^11,12^. Based on a sequence analysis, the N-terminal half of the LRRC8 isoforms is predicted to form four transmembrane (TM) helices, while the C-terminal half of the LRRC8 isoforms is predicted to contain up to 17 leucine-rich repeats (LRR). A previous bioinformatics analysis suggested that the N-terminal halves of the LRRC8 isoforms share weak homology to pannexin, connexin, and innexin^13^. Since the previous structural analyses of the gap junction channels, connexin and innexin, revealed that their hemichannels form a hexamer and an octamer, respectively^14–16^, the LRRC8 proteins are also predicted to form hexameric or higher TM region oligomers to create the channel pore^13^. Further electrophysiological analyses, using cells expressing LRRC8 isoforms and LRRC8 isoforms embedded in lipid bilayers, demonstrated that the heteromeric LRRC8 complex, consisting of LRRC8A and at least one of the four other LRRC8 isoforms, is indispensable for the channel activity in cells, while the homomeric LRRC8A protein still retains the hypotonically induced channel activity under lipid embedded conditions^10,17^. This heteromeric assembly of LRRC8A and the other LRRC8 isoforms contributes to the diversity of the channel properties, including the gating kinetics and modulation, as well as the substrate specificity. For example, cells expressing the LRRC8A/C heteromer are inactivated more slowly than those expressing the LRRC8A/E heteromer^10^. In addition, the activity of the LRRC8A/E heteromer is potentiated by intracellular oxidation, while the activities of the LRRC8A/C and LRRC8A/D heteromers are inhibited by intracellular oxidation^18^. Moreover, cells expressing the LRRC8A/D heteromer release uncharged osmolytes such as taurine, and uptake anti-cancer drugs, including cisplatin, while those expressing the LRRC8A/C/E heteromer release charged organic osmolytes^19–21^.

As the overall sequence of LRRC8 has little similarity to any other known classes of ion channels, the detailed structural information of the LRRC8 protein has long been awaited to elucidate the molecular architecture of LRRC8 and to clarify the molecular mechanisms, such as substrate permeability. Here, we report the cryo-electron microscopy (cryo-EM) structure of the homomeric human LRRC8A, at an average resolution of 4.25 Å (local resolution of the TM region is ∼3.8 Å). The present structure provides a framework for understanding the mechanisms of this unique class of ion channels.

## Results

### Structure determination of homomeric HsLRRC8A

We first screened the LRRC8 proteins from various vertebrates by Fluorescence-detection Size-Exclusion Chromatography (FSEC) and an FSEC-based thermostability assay (FSEC-TS)^22,23^, and found that the human LRRC8A protein (HsLRRC8A; the “Hs” refers to *Homo sapiens*), the best characterized protein among the known LRRC8 proteins, produced symmetric and monodisperse FSEC peaks when solubilized with digitonin (Extended Data Fig. 1a). We also confirmed the hypotonicity-induced channel activity of the purified HsLRRC8A protein in reconstituted liposomes (Extended Data Fig. 1b,c). Although the LRRC8 proteins reportedly function as heteromers in cells^10^, the structure of the homomeric HsLRRC8A protein may provide important insights into the basic architecture of the LRRC8 proteins, given the sequence similarity between the isoforms (Extended Data Fig. 2). Therefore, we set out to elucidate the structure of the homomeric HsLRRC8A. The purified HsLRRC8A protein was vitrified on grids, and images of the grids were recorded by electron microscopy (Extended Data Fig. 3a). In the 3D classification step, classes #1 and #2 showed clearer densities and similar assemblies, respectively, as compared to the other classes. Therefore, we chose these two classes for further refinement (Extended Data Fig. 3b,c). Since these classes exhibited 3-fold rotational symmetry, C3 symmetry was imposed in the subsequent reconstructions. In the final 3D map reconstructed to an overall 4.25 Å resolution (local resolution of the TM region is ∼3.8 Å) (Extended Data Fig. 3d,e), the densities of the six protomers of HsLRRC8A (namely, the α to ζ subunits) were clearly observed, including the long arc-like densities corresponding to the leucine-rich repeat (LRR) region. This density map enabled us to assign the topologies of the LRR and other regions, including the extracellular, transmembrane (TM), and intracellular regions (Fig. 1a-c, Extended Data Fig. 3b, c). The local resolution of the core of the TM region is estimated to be ∼3.8 Å, and its density is sufficiently resolved to enable the tracing of the α-helices and the bulky amino-acid side chains (Fig. 1a, b). The densities corresponding to most of the extracellular and intracellular regions are also resolved well enough to trace the secondary structures, using the structure of the previously determined nematode Innexin 6 (CeInnexin-6; the “Ce” refers to *Caenorhabditis elegans*) (PDB ID: 5H1Q) as a guide. Two loops connecting the TM helices in these regions are disordered (Fig. 1d, e). In contrast to the clear densities of the extracellular, TM and intracellular regions, the local resolution of the LRR regions is around 5.5 Å, and only the main chain of the LRR region is resolved in the density map (Fig. 1a, b). Therefore, we generated the homology model of the LRR region of HsLRRC8A based on the previously determined LRR protein structure (PDB ID: 4U08) using the Phyre2 server^24^, and used it to guide the model building. The final model of HsLRRC8A, in which about 90% of the residues were assigned, contains the residues R18-D806 and is missing parts of the extracellular loop (T61-D91) and the intracellular loop (T176-R226 in the α, γ, and ε subunits; T176-L232 in the β, δ, and ζ subunits).

**Figure 1.**
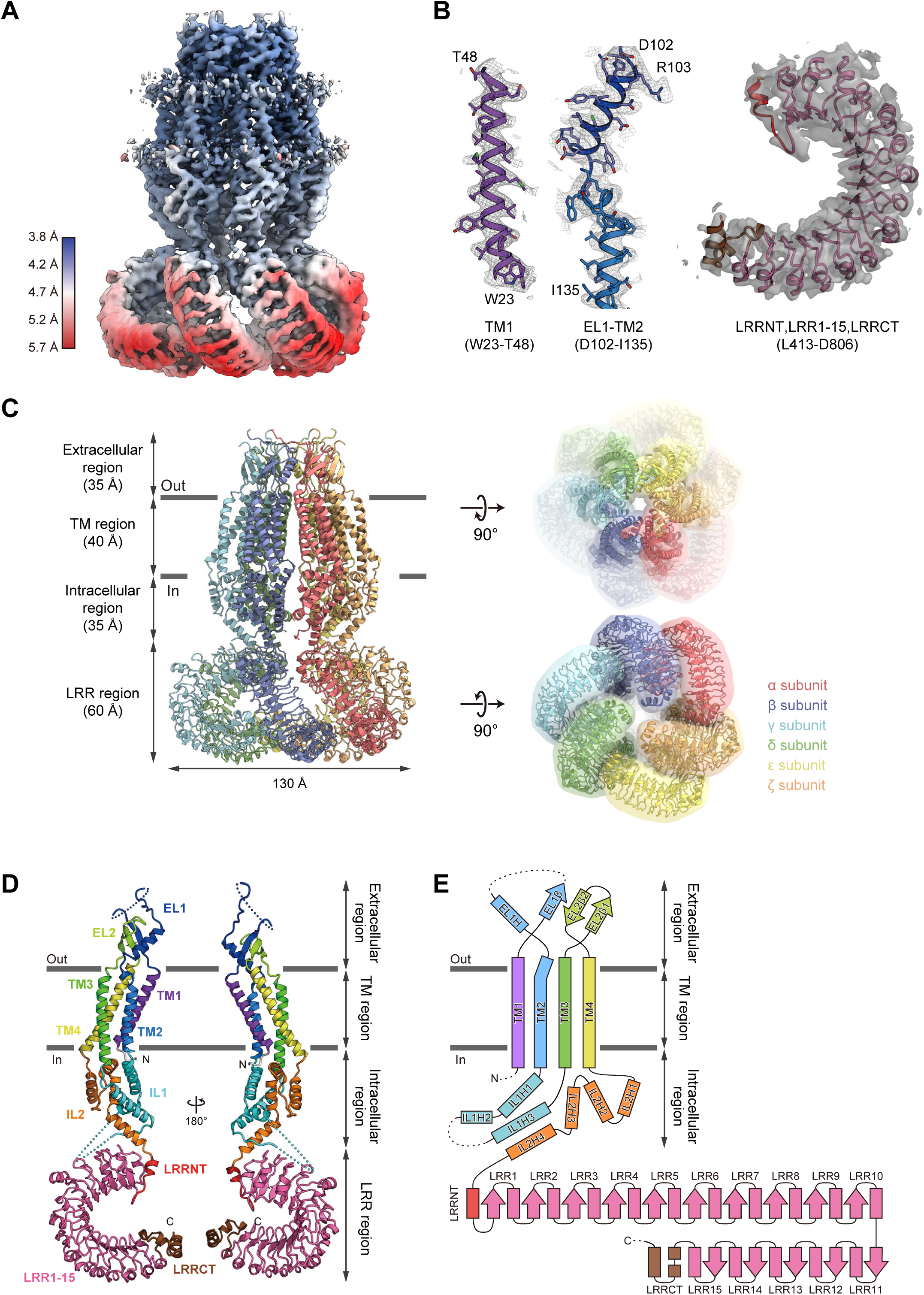
Overall structure and subunit interactions of the HsLRRC8A hexamer. (**a**) Local resolution of the HsLRRC8A structure, estimated by RELION. (**b**) Representative density maps of the transmembrane and LRR regions. The cartoon models of the TM1, EL1 and the following TM2, and LRR regions were fitted to the density maps (colored gray). (**c**) Overall structure of the HsLRRC8A hexamer, viewed parallel to the membrane (left). Overall structure of the HsLRRC8A hexamer with molecular envelope, viewed from the extracellular side (right, upper) and the intracellular side (right, lower). (**d**) The subunit structures of HsLRRC8A, viewed parallel to the membrane. Each region is colored as follows: TM1, purple; EL1, blue; TM2, light blue; IL1, cyan; TM3, green; EL2, light green; TM4, yellow; IL2, orange; LRRNT, red; LRR1-15, pink; and LRRCT, brown. The N-and C-termini are indicated by ‘N’ and ‘C’, respectively. (**e**) Schematic representation of the HsLRRC8A membrane topology, colored as in (d). The dotted lines indicate the disordered linkers. The molecular graphics were illustrated with CueMol (http://www.cuemol.org/).

### Overall structure and subunit fold of HsLRRC8A

The hexameric structure of HsLRRC8A adopts an upside-down sea anemone-like shape, when viewed parallel to the membrane (Fig. 1c). The subunit structure consists of four regions; *i.e*., the extracellular region (Q49-H119 and H287-L314), TM region (W23-T48, W120-F144, I259-V286, and A315-R345), intracellular region (K145-D258 and R346-T412), and LRR region (L413-D806) (Fig. 1c-e), with both the N-and C-termini residing on the cytoplasmic side. The overall structures of the six subunits are similar, except for several points as described later, and thus we mainly discuss the structure of the α subunit of the hexamer. The extracellular region protrudes by ∼35Å above the cell membrane, and forms the channel pore with the TM region. The LRR region is protruded toward the intracellular side, and is twisted in a clockwise manner as viewed from the cytoplasmic side (Fig. 1c). The TM region consists of the four TM helices (TM1-4), which are connected by the two extracellular loops (EL1 and EL2) of the extracellular region, and the intracellular loop (IL1) of the intracellular region (Fig. 1d, e). IL2 in the intracellular region connects the TM and LRR regions (Fig. 1d, e). EL1 possesses one α-helix (EL1H) and one β-strand (EL1β), while EL2 possesses two β-strands (EL2β1 and EL2β2) (Fig. 1d, e). IL1 possesses three α-helices (IL1H1–3), while IL2 possesses four α-helices (IL2H1–4) (Fig. 1d, e). EL1 and IL1 contain disordered regions (T61–D91 and T176–R226), suggesting that these loops have conformational flexibility (Fig. 1d, e, Extended Data Fig. 2). While a previous bioinformatics analysis suggested that LRRC8 contains a 17 LRR repeat^12^, the present structure revealed that the LRR region consists of the leucine-rich repeat N-terminal helix (LRRNT), fifteen leucine-rich repeats (LRR1–15), and the following C-terminal helices (LRRCT) (Fig. 1d, e, Extended Data Fig. 2).

### Differences in structures and interactions among HsLRRC8A subunits

With its global C3 symmetry, the hexameric HsLRRC8A structure exhibits the “trimer of dimers” architecture; in other words, the structures of the α, γ, and ε subunits, and those of the β, δ, and ζ subunits, are identical (Fig. 1c, 2a). In contrast, a distinctive structural difference is observed between the α and β subunits. The LRR region is rotated by ∼22° relative to the extracellular, TM, and intracellular regions (Fig. 2b). No large structural difference is observed in each region, as their structures superimpose well with RMSD values less than 1 Å (0.39 Å, 0.40 Å, 0.98 Å, and 0.47 Å for the extracellular, TM, intracellular, and LRR regions, respectively), suggesting their structural rigidity (Fig. 2b, Extended Data Fig. 4). Therefore, this structural difference between the subunits can be ascribed to the rotation of the IL2H4 helix in the IL2 loop, which connects the intracellular and LRR regions as a hinge (Fig. 2c).

As a result of this structural difference between the subunits, several variations are observed in the interfaces between the subunits; *e.g*., the α subunit tightly interacts with the β subunit, while it loosely interacts with the ζ subunit (Fig. 2a). Notably, in the “tight” interaction, there are extensive contacts between the adjacent subunits from the extracellular region to the LRR region, whereas in the “loose” interaction, there are almost no contacts between the adjacent subunits in the lower part of the TM region, the intracellular region, and the upper part of the LRR region (Fig. 2a). At the TM region, the subunit interface is highly hydrophobic and mainly formed by the TM2 helix from one subunit, and the TM1 and TM4 helices from the other adjacent subunit (Extended Data Fig. 5a, b). There is a “lateral crevice” open to the lipid bilayer across the central channel pore, and it is wider in the loose interface than in the tight interface (Extended Data Fig. 5a, b). At the intracellular region, the subunit interface is highly hydrophilic and formed by the IL1H1 and IL1H3 helices from one subunit, and the IL1H2, IL2H2 and IL2H3 helices from the other adjacent subunit (Extended Data Fig. 5c, d). In this region, extensive interactions are formed in the tight interface, while few interactions are observed in the loose interface (Extended Data Fig. 5c, d). At the LRR region, extensive interactions throughout all of the LRR repeats are observed in the tight interface, while in the loose interface, fewer interactions are formed between the C-terminal LRRCT helices from one subunit and the middle of the LRR repeats from the other adjacent subunit (Extended Data Fig. 5e, f). Notably, a heterozygous LRRC8A truncation mutant, in which the C-terminal 91 amino acids are replaced by 35 amino acids encoded in the intron sequences, has been reported to suppress the LRRC8A function, which causes an innate immune disease^8,9^. This disease-related truncation might abrogate the interactions at this loose interface, thereby affecting the proper formation of the hexameric channel structure. In contrast, the interactions in the tight and loose interfaces of the extracellular region are almost the same (Extended Data Fig. 5g, h). The interactions are formed by the extensive contacts between the EL1H helix and the three-stranded β-sheet (EL1β, EL2β1, and EL2β2) from the other adjacent subunit (Extended Data Fig. 5g, h).

**Figure 2.**
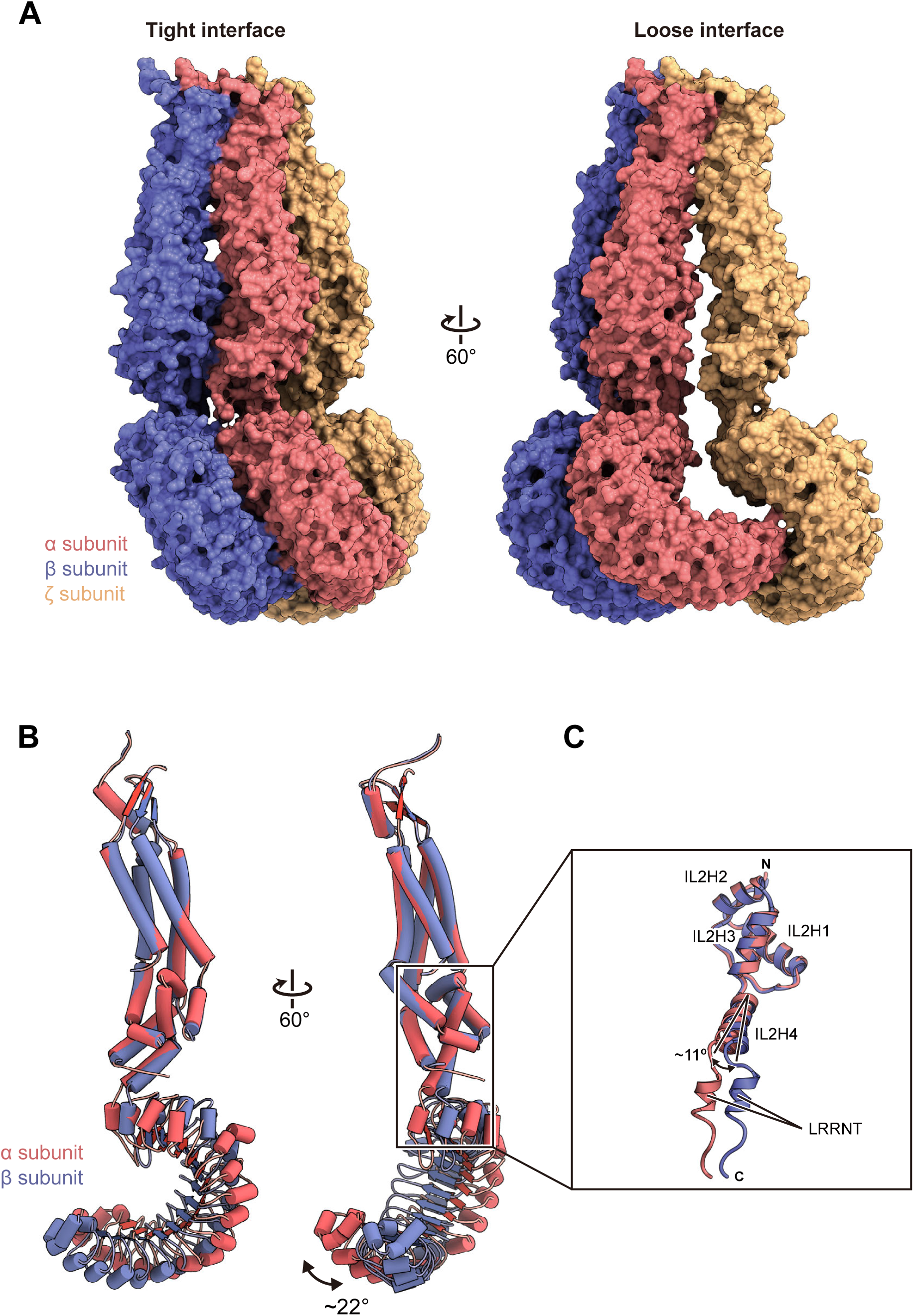
Subunit Interface. (**a**) The tight and loose subunit interfaces (left and right panels, respectively) between the three neighboring subunits (α, β, and ζ) of HsLRRC8A. The structures of the subunits are shown as the solvent-accessible surface models. (**b**) Two adjacent subunits (α and β subunits) of HsLRRC8A are superimposed on each other, using only the TM region. The RMSD value for the 553 Cα atoms from the subunits is 3.79 Å. (**c**) Close-up view of the superimposed intracellular IL2 regions from two adjacent subunits. The residues involved in helix formation are depicted in stick representations. The RMSD value for the 66 Cα atoms from the subunits of the IL2 regions is 0.88 Å.

Overall, these structural comparisons between the subunits suggest that the flexibility around the IL2H4 helix, which connects the intracellular and LRR regions, causes the structural differences between the subunits, which may enable the formation of the “trimer of dimers” architecture of the hexameric channel, with the wide crevice in the loose interface.

### Channel pore

HsLRRC8A has a channel pore along the central axis, perpendicular to the membrane. The channel pore is mainly composed of the EL1H helix in the extracellular region and the TM1 helix in the TM region, as well as the IL1H1 and IL1H3 helices in the intracellular region (Fig. 3a). The diameter of the pore is about ∼ 25 Å on the intracellular side, and it widens to about 50 Å at the lower part of the TM region, and then narrows to less than 10 Å at the upper part of the extracellular region (Fig. 3b-e). Therefore, the channel pore formed by the hexameric LRRC8A is accessible from both the extracellular and intracellular sides, and thus the present structure of LRRC8A probably represents the open conformation. The residues lining the channel pore are mainly hydrophilic, suggesting that the pore is filled with water molecules in the present structure. A previous experiment revealed that the T44C mutation alters the I^−^ versus Cl^−^ anion selectivity of HsLRRC8A, and the MTSES (2-sulfonatoethyl methanethiosulfonate) modification of the T44C mutant significantly reduced the channel activity, suggesting that T44 is involved in the pore formation^9^. Consistently, in our HsLRRC8A structure, T44 is located on TM1 and involved in the formation of the solvent-accessible surface (Fig. 3b, d). The most-constricted site of the pore has a diameter of about 7.6 Å, which is wide enough to permeate a hydrated Cl^−^ ion. This constriction site is mainly formed by the side chains of the positively charged R103 residues (Fig. 3b, c). In addition, a negatively charged residue, D102, is located next to R103, and is positioned just above R103 like a “lid” for the constriction site. These structural features suggest that the D102 and R103 residues play an important role in the ion permeation (Fig. 3b, c). Furthermore, a recent electrophysiological analysis demonstrated that three residues, corresponding to K98, Y99, and D100 in the LRRC8A isoform, are involved in the voltage dependence and kinetics of the LRRC8 isoforms^25^. Single and combination mutants of these residues reportedly affected the rate and voltage dependence of inactivation, the kinetics of inactivation, and the ion selectivity^25^. Consistently, these three residues are located in the loop preceding the EL1H helix and are exposed to the extracellular entrance of the channel pore in the present HsLRRC8A structure (Fig. 4a), suggesting their importance for the ion permeation.

**Figure 3.**
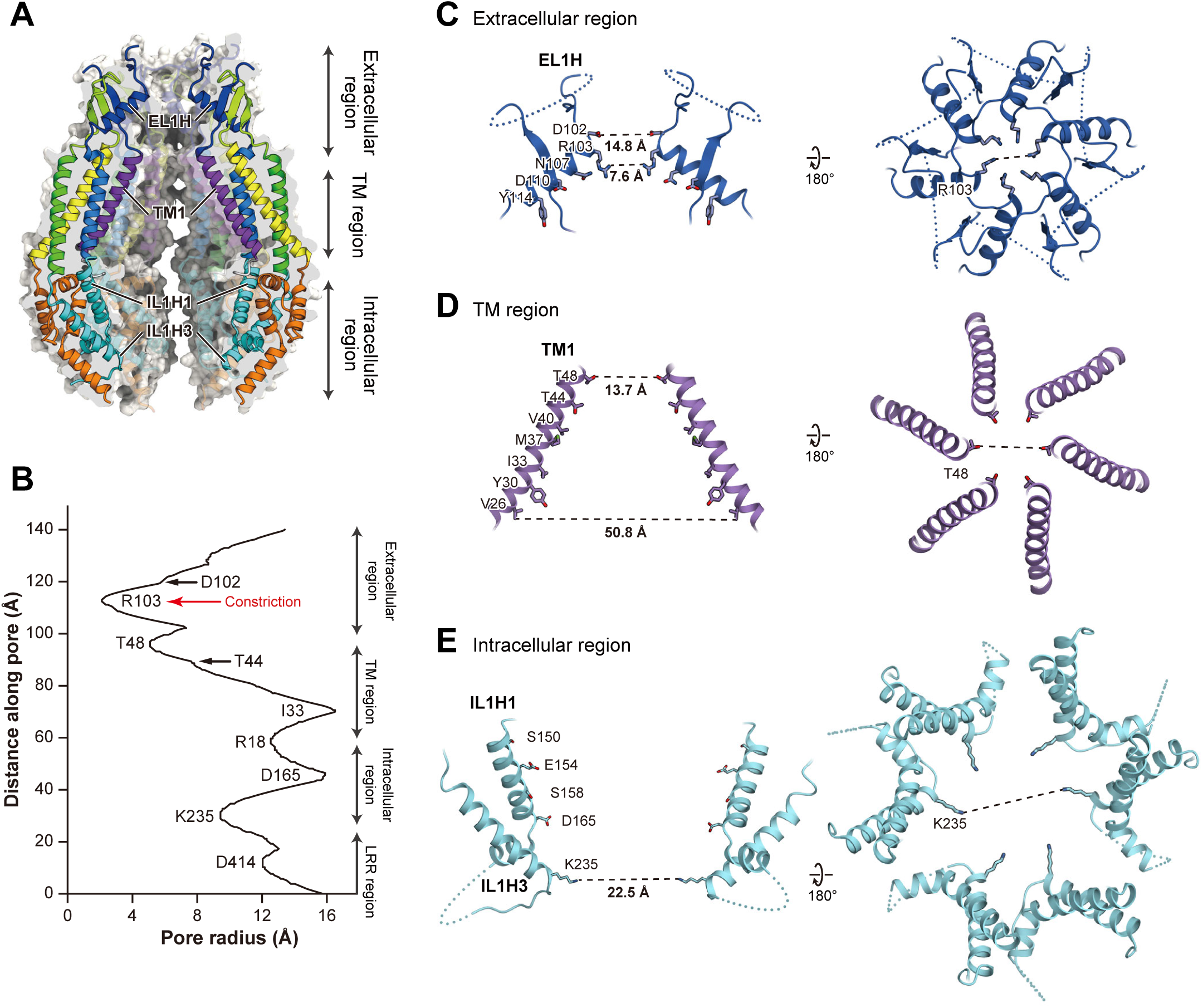
Ion channel pore. (**a**) Cross-sections of the surface representation for the N-terminal half of the HsLRRC8A subunits, showing the ion channel pore. The four subunits are colored according to Fig. 1d, and are shown in cartoon representations. (**b**) The pore radius for the HsLRRC8A structure along the pore center axis. The pore size was calculated with the program HOLE^40^. (**c-e**) Close-up views of the channel pore forming regions. In the left views, only two diagonal subunits viewed parallel to the membrane are shown, for clarity. The side chains of the pore-lining amino residues are depicted in stick representations. The distances between the residues involved in the pore constrictions are shown. In the right views, all six subunits viewed from the extracellular side are shown.

**Figure 4.**
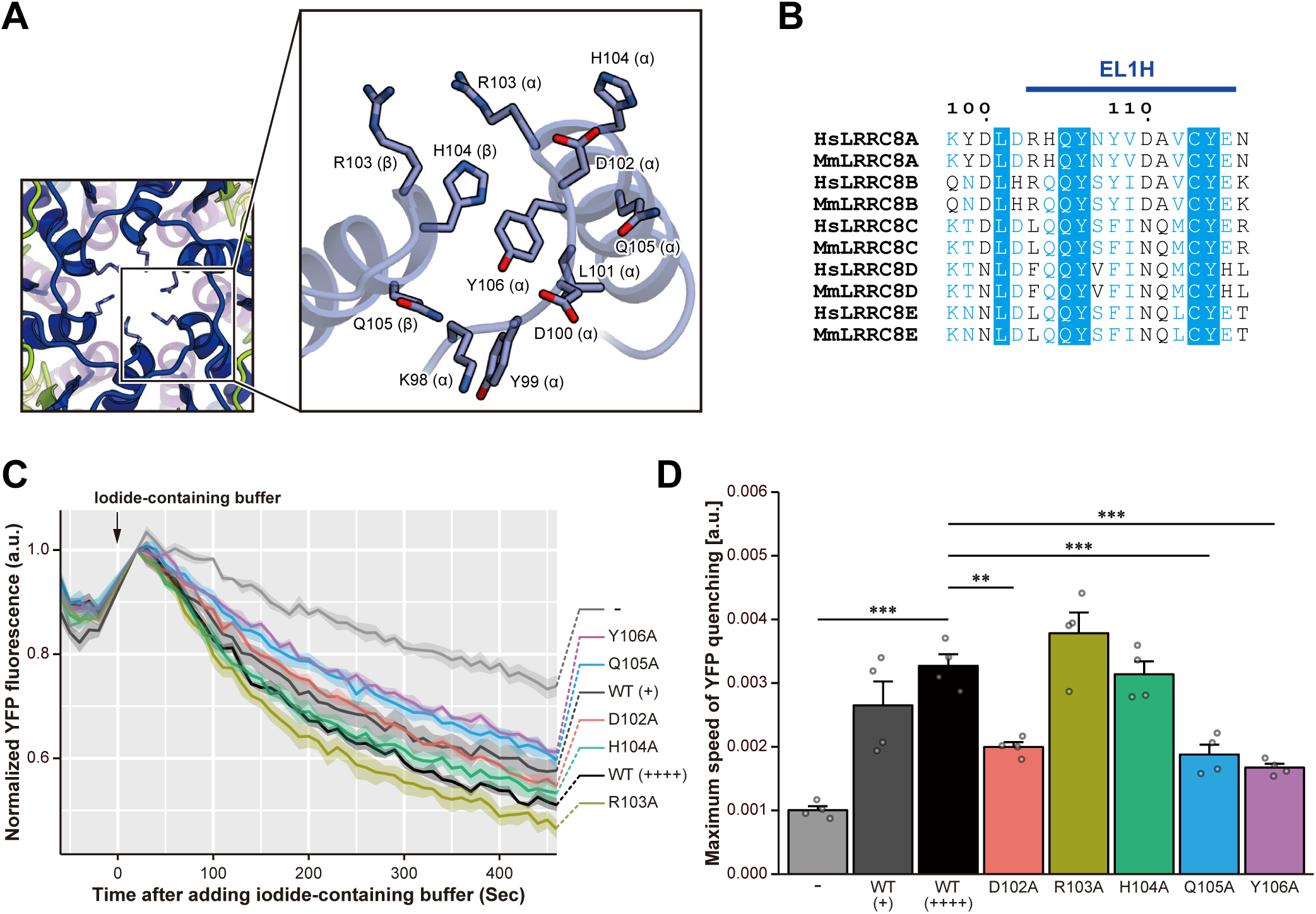
Pore constriction site. (**a**) Close-up view of the neighboring EL1 helices forming the pore constriction site. The side chains are depicted by stick models. (**b**) Sequence alignment around the pore constriction site, depicted according to Extended Data Fig. 2. (**c**) VRAC activity in the LRRC8A-rescued *LRRC8A*-knockout HEK293A cells. Each line indicates the mean ± s.e.m. of the normalized YFP fluorescence (*n* = 4). Hypotonicity after the addition of iodide: 231 mOsm. WT (+) and WT (++++) represent the different expression levels of the LRRC8A wild-type protein detected by western blotting, according to Extended Data Fig. 6c. (**d**) Maximum speed of hypotonicity-induced YFP quenching. Data are presented as the mean ± s.e.m. with each individual light grey point (n = 4). ** P < 0.01, *** P < 0.001 by Dunnett’s test.

Based on the present HsLRRC8A structure, as well as the previous functional analysis, we hypothesized that the N-terminal tip of the EL1H helix (D102-Y106) plays a critical role in the ion permeation, as it resides on the extracellular side of the channel pore and constitutes its most-constricted site (Fig. 4a,b). To corroborate this notion, we generated a series of single alanine mutants of residues D102 to Y106 in HsLRRC8A and performed cell-based measurements of the VRAC activity, using assays originally developed upon the discovery of the LRRC8A protein as a VRAC component^9,10^. First, by a confocal microscopic analysis, we confirmed that the signals derived from these mutants are observed at the cell membrane (Extended Data Fig. 6a). In addition, the FSEC analysis of these mutants produced symmetric and monodisperse peaks, similar to those of the wild-type channel, confirming the proper structural integrities of these mutant channels (Extended Data Fig. 6b). We transfected the halide-sensitive YFP mutant^26^ with or without HsLRRC8A wild-type (WT) into HEK293A cells, in which the endogenous *LRRC8A* gene is knocked out, and verified the assay system: HsLRRC8A WT could rescue the hypotonicity-induced iodide quenching response in the *LRRC8A* KO cells (Fig. 4c, d). We then evaluated each of the HsLRRC8A mutants within the range of WT expression levels (Extended Data Fig. 6c), and found that the D102A, Q105A and Y106A mutants decreased the quenching speed as compared to the wild-type channel (Fig. 4c, d, Extended Data Fig. 6d). Even though the reduction of the quenching speed cannot exactly explain how the mutations affect the channel properties, such as the rate of ion permeation and the channel open probability, the results support our hypothesis that the N-terminal tip of the EL1H helix is important for the ion permeation. Interestingly, the amino-acid sequences in this region of the other LRRC8 isoforms deviate from that of LRRC8A (Fig. 4b). These differences in the amino acid sequences in this region may alter the properties of this constriction site in the channel pore, thereby modulating the substrate specificity of the heteromeric LRRC8 hexamers containing different isoforms^20,21^.

### Putative structural change of LRRC8

During the 3D classification, in addition to classes #1 and #2 used for the 4.25 Å reconstruction, we found that class #7 also showed clearer LRR densities than the other classes (Extended Data Fig. 3b). Interestingly, class #7 exhibited the distorted arrangement of the LRR densities, as compared to those in classes #1 and #2, and thus it may represent another structural population of LRRC8A. Therefore, we performed the 3D reconstruction of class #7 without imposing any symmetry. The overall resolution of the resulting map was 9.17 Å, and we could assign the extracellular, TM, intracellular, and LRR regions. We then constructed a model of class #7, based on the higher resolution structure.

In this model constructed from class #7, the LRR region is substantially deviated and more loosely packed as compared to that in the high-resolution structure from classes #1 and #2 (Fig. 5a, b). Accordingly, we refer to this class #7 structure as the “relaxed form” (Fig. 5a), and the higher-resolution structure from classes #1 and #2 as the “compact form” (Fig. 5b). In the relaxed form, the LRR regions are further tilted and expanded away from the central axis, with their width elongated by ∼20 Å and height shortened by ∼8 Å, as compared with the compact form (Fig. 5a, b). The C3 symmetry observed in the compact form is absent and the arrangement of the LRR regions is completely asymmetric. In contrast, the arrangement of the secondary structure elements in the extracellular, TM, and intracellular regions is quite similar to that observed in the compact form. The structures of the LRR regions in both the relaxed and compact forms were also similar, including their curvatures. Thus, the overall structural difference between the relaxed and compact forms can be explained by the rigid-body motions of the LRR and the other regions, in which the IL2H4 helix in the IL2 loop serves as a hinge. Interestingly, a similar structural difference is also observed between the α and β subunits in the compact form (Fig. 2, Extended Data Fig. 4). These structural differences suggest the structural flexibility around the IL2H4 helix in the IL2 loop, which might be involved in the important structural transition for the channel function (Fig. 5c).

**Figure 5.**
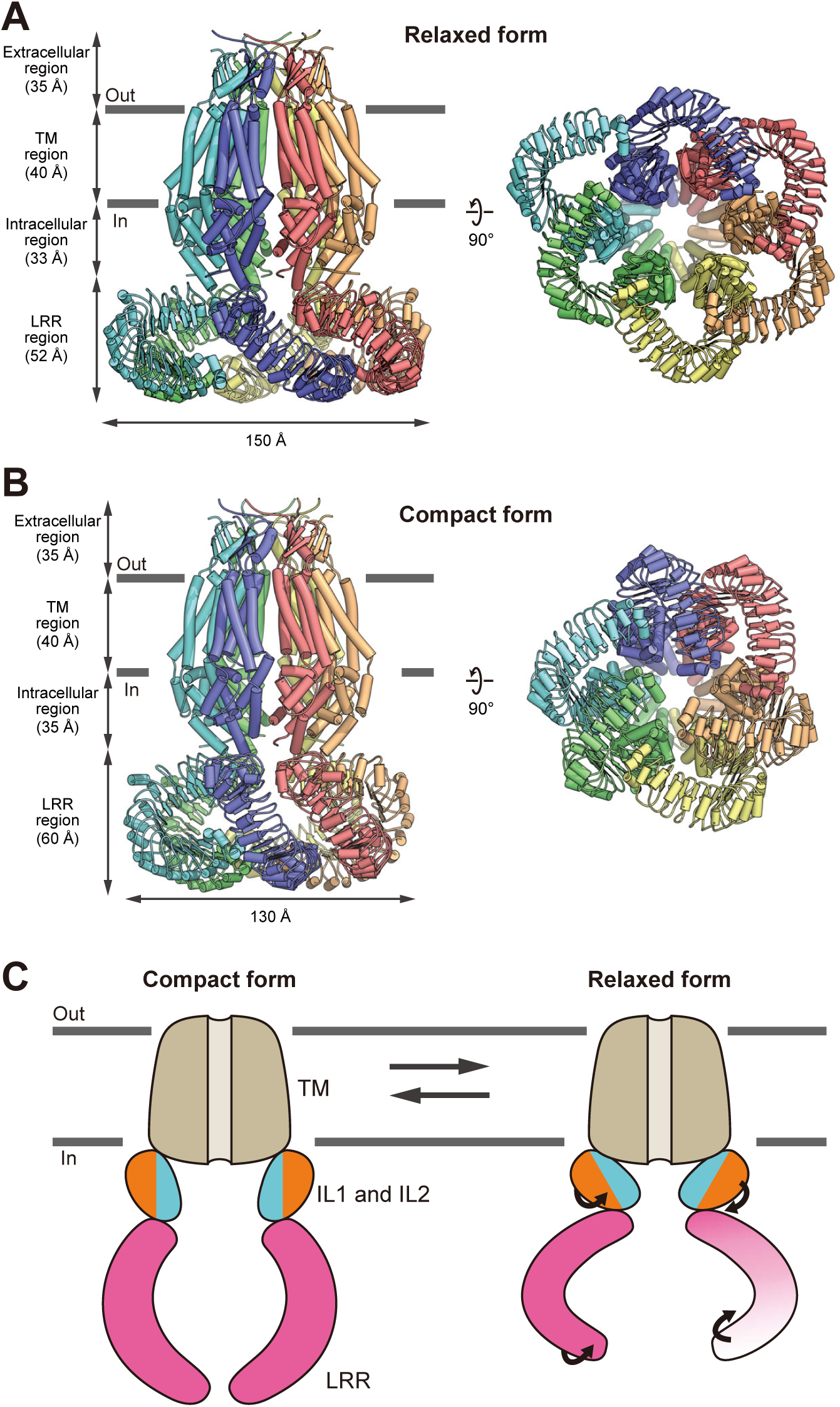
Compact and relaxed structures of HsLRRC8A. (**a,b**) Overall structure of the HsLRRC8A hexamer in the relaxed form (a), and in the high-resolution compact form (b), viewed parallel to the membrane (left), and from the extracellular side (right). The structures are depicted in cylinder representations. (**c**) Schematic drawing depicting the structural transition between the compact and relaxed forms of HsLRRC8A.

### Structural comparison with gap junction channels

Consistent with the previous bioinformatics analysis^13^, the extracellular, TM, and intracellular regions of HsLRRC8A share several structural features with those of the CeInnexin-6 and human connexin 26 (HsConnexin-26) gap junction channels (Extended Data Fig. 7)^14,15^, although they have low sequence identity (∼15% with CeInnexin-6; ∼18% with HsConnexin-26). In the TM region, they share the same topology of four TM helices, in which TM1 is the innermost helix. In the extracellular region, they commonly possess conserved disulfide bonds between the EL1 and EL2 loops, and common secondary structure elements, including helix EL1H, as well as the β-sheet bridging the EL1 and EL2 loops (Extended Data Fig. 7a-c). In the intracellular region, helices IL1H1 and IL1H3 in the IL1 loop are also conserved in CeInnexin-6 (Extended Data Fig. 7a, b). In contrast, HsLRRC8A possesses several distinct structural features as compared to the gap junction channels. First, the subunit interfaces of HsConnexin-26 and CeInnexin-6 are tightly packed and the large crevice is not observed (Extended Data Fig. 7d-f)^14,15^, suggesting that the crevice of LRRC8 is involved in its specific function. Second, the HsConnexin-26 and CeInnexin-6 structures revealed that their short N-terminal extension (∼15 residues) forms a short α helix (NTH), which is involved in the channel pore formation (Extended Data Fig. 7g-i). This NTH was speculated to be important for the ion permeation and gating in the gap junction channels^14,15^. HsLRRC8A also possesses a short extension at its N terminus; however, its density was not observed in the present cryo-EM structure. The amino-acid sequence of the N-terminal extension of HsLRRC8A is quite different from those of both HsConnexin-26 and CeInnexin-6, and is not predicted to form a helix, unlike the cases of HsConnexin-26 and CeInnexin-6 (Extended Data Fig. 7j). Thus, the N-terminal extension of LRRC8 would not form a helix and is likely to play a different role from those of the gap junction channels. Third, in the TM region, while TM1 in HsLRRC8A is straight and faces toward the intracellular side, those in the gap junction channels are bent at the middle, and consequently the N-termini of the gap junction channels face toward the extracellular side to form the channel pore (Extended Data Fig. 7b, c). Fourth, in the intracellular region, the IL1 and IL2 loops of HsLRRC8A (144 residues in IL1; 67 residues in IL2) are longer than those of CeInnexin-6 (54 residues in IL1; 59 residues in IL2) (Extended Data Fig. 7a, b). As a result, in HsLRRC8A, the long linker between the IL1H2 and IL1H3 helices in IL1, as well as the IL2H4 helix in IL2, directly interacts with the LRR region. Particularly, the IL2H4 helix, which connects the intracellular and LRR regions, only exists in the LRRC8 isoforms. Fifth, in the extracellular region, EL1H in HsLRRC8A (14 residues) is longer than those in the gap junction channels (10 residues), and the N-terminal tip of the EL1H helix in HsLRRC8A forms the channel pore with the TM1 helix (Extended Data Fig. 7a-c, g-i). In addition, the sizes and structures of the EL1 and EL2 loops of the gap junction channels are different from those of HsLRRC8A (Extended Data Fig. 7a-c). Indeed, the extracellular regions of HsConnexin-26 and CeInnexin-6 were shown to interact with each other to form the gap junction channel, in which the EL1 and EL2 loops provide the interaction interface^14,15^. Therefore, these structural differences in the subunit interfaces, as well as the extracellular, TM, and intracellular regions, contribute to the functional diversity between the volume-regulated channels and the gap junction channels.

## Discussion

In this study, we present the homomeric human LRRC8A cryo-EM structure, the first structure of a member of the LRRC8 protein family responsible for the VRAC function. The overall structure, reconstructed at 4.25 Å resolution (∼3.8 Å in the TM region), exhibited a hexameric assembly with a trimer of dimers, rather than a symmetrical hexamer, revealing the architecture of this unique class of ion channels (Fig. 1) The channel pore observed along the central axis is open toward both the extra-and intracellular sides, suggesting that the present structure represents the channel-open form (Fig. 3).

Although LRRC8A is an essential component of the VRAC activity, overexpression of the LRRC8A gene in the quintuple-knockout LRRC8^-/-^ cells or quadruple-knockout LRRC8(B/C/D/E)^-/-^ cells did not rescue the VRAC activity, suggesting that the homomer of LRRC8A does not form the active VRAC pore *in vivo*. Nevertheless, interestingly, the homomer of LRRC8A reconstituted into lipid droplet bilayers still retains VRAC characteristics, including hypotonicity-induced currents and sensitivity to a specific VRAC inhibitor, DCPIB (4-(2-butyl-6,7-dichlor-2-cyclopentyl-indan-1-on-5-yl) oxybutyric acid)^17^. We also confirmed that the homomeric HsLRRC8A protein used in this study has hypotonicity-induced channel activity upon osmotic change, using the reconstituted liposomes (Extended Data Fig. 1b,c). Furthermore, a previous biochemical experiment showed that the overall assembly, including the oligomeric state, of the LRRC8A homomer is similar to those of the LRRC8A-containing heteromers. The present structure of the homomeric HsLRRC8A provides important structural insights into the LRRC8A-containing heteromers. To visualize the conserved and diversified sites among the LRRC8 gene family members, we calculated the conservation score using the program ConSurf^27^, and mapped the score on the molecular surface of HsLRRC8A (Extended Data Fig. 8a). The result showed that the subunit interface and the channel pore in the TM, extracellular, and intracellular regions, except for the constriction site, are highly conserved among the LRRC8 gene family members. In contrast, the membrane side of the TM region, the overall LRR region, and the constriction site of the pore exhibited relatively low conservation scores. Next, based on the present structure of HsLRRC8A, we constructed a model structure of the HsLRRC8 heteromer containing HsLRRC8D (54% sequence identity with HsLLRC8A; Extended Data Fig. 8b). Although the stoichiometry of the heteromeric LRRC8 is still unknown, this (LRRC8A)_5_·LRRC8D heteromer may be one of the oligomeric states present in the cellular context^17^. In this model structure, no large steric clashes are observed at the subunit interfaces (Extended Data Fig. 8b), suggesting that the formation of the hexamer, as well as the crevice between the subunits, is plausible in the LRRC8A-containing heteromers. In contrast, the structure of the pore constriction site, which is mainly formed by the helix EL1H, is different from that of the homomeric LRRC8A (Extended Data Fig. 8b). These results again underscore the notion that the sequence diversity in the pore constriction site is important for the functional diversity of the LRRC8A-containing heteromers.

In the subunit interface of the TM region, we observed the lateral crevice formed between the subunits, which is opened toward the membrane side (Fig. 2a). Although it is currently unknown whether the activity of LRRC8 proteins is influenced by the lipid environment, it is possible that the LRRC8 proteins might directly interact with the lipid molecules through these crevices, in a similar manner to other channels, such as the mechanosensitive two-pore domain K^+^ channels^28^. A recent electrophysiological analysis using purified LRRC8 proteins demonstrated that low ionic strength alone is sufficient for their activation, and it was speculated that the sensor for the ionic strength exists in the IL1 loop, as well as in LRRCT^17^. Indeed, the long linker between the IL1H2 and IL1H3 helices, which is largely disordered in the present structure, contains several charged residues that are conserved among LRRC8 isoforms (Extended Data Fig. 2). Although the question of whether the ionic strength directly modulates the channel activities of the LRRC8 proteins remains to be answered, these observations lead us to speculate that the interaction of the IL1 loop with ions affects its conformation, thereby changing the arrangement of the LRR and other regions, as observed in the present relaxed form (Fig. 5a, c), to regulate the channel activity. To understand the gating mechanism of the LRRC8 protein depending on the cellular volume change, the structural analysis of the closed form of LRRC8 is required.

## Methods

### Protein expression, purification and sample preparation for cyro-EM

The human LRRC8A WT protein (NCBI Reference sequence number: NP_062540.2) was cloned from human brain cDNA (ZYAGEN) into the pEG BacMam vector, with an C-terminal GFP-His8 tag and a tobacco etch virus (TEV) cleavage site, and was expressed in HEK293S GnTI^−^ (N-acetylglucosaminyl-transferase I-negative) cells (ATCC, cat. no. CRL-3022)^29^. Cells were collected by centrifugation (6000×g, 10 min, 4 °C) and disrupted by sonication in buffer (50 mM Tris, pH 8.0, 150 mM NaCl) supplemented with 5.2 µg/ml aprotinin, 2 µg/ml leupeptin, and 1.4 µg/ml pepstatinA (all from Calbiochem). Cell debris was removed by centrifugation (10,000×g, 10 min, 4 °C). The membrane fraction was collected by ultracentrifugation (138,000×g, 1 h, 4 °C). The membrane fraction was solubilized for 1 h at 4 °C in buffer (50 mM Tris, pH 8.0, 150 mM NaCl, 5 mM dithiothreitol (DTT), 1% digitonin (Calbiochem)). Insoluble materials were removed by ultracentrifugation (138,000×g, 1 h, 4 °C). The detergent-soluble fraction was incubated with CNBr-Activated Sepharose 4 Fast Flow beads (GE Healthcare) coupled with an anti-GFP nanobody (GFP enhancer)^30^, and incubated for 2 h at 4 °C. The beads were washed with SEC buffer (50 mM Tris, pH 8.0, 150 mM NaCl, 5 mM DTT, 0.1% digitonin), and further incubated overnight with TEV protease to remove the GFP-His8 tag. After TEV protease digestion, the flow-through was collected, concentrated, and purified by size-exclusion chromatography on a Superose 6 Increase 10/300 GL column (GE Healthcare), equilibrated with SEC buffer. The peak fractions of the protein were collected and concentrated to 15 mg/ml, using a centrifugal filter unit (Merck Millipore, 100 kDa molecular weight cutoff). A 3 µl portion of the concentrated HsLRRC8A protein (15 mg/ml) was applied to a glow-discharged Quantifoil R1.2/1.3 Cu/Rh 300 mesh grid (Quantifoil), blotted using Vitrobot Mark IV (FEI) under 4°C and 100% humidity conditions, and then frozen in liquid ethane.

### EM image acquisition and data processing

The grid images were obtained with a Tecnai Arctica transmission electron microscope (FEI) operated at 200 kV, and recorded by a K2 Summit direct electron detector (Gatan) operated in the super-resolution mode with a binned pixel size of 1.490 Å. The dataset was acquired with the SerialEM software^31^. Each image was dose-fractionated to 40 frames at a dose rate of 6-8 e^−^/pixel/sec, to accumulate a total dose of ∼50 e^−^/Å^2^. In total, 5,305 super-resolution movies were collected. The movie frames were aligned in 5 × 5 patches, dose weighted and binned by two in MotionCor2 (ref. 32), and down-sampled by two. Defocus parameters were estimated by CTFFIND 4.1 (ref. 33). First, template-based auto-picking was performed with 2D class averages of a few hundred manually picked particles as templates^34^. A total of 1,421,314 particles were extracted in 3.25 Å/pix. These particles were divided into four batches and subjected to three rounds of 2D classification in RELION 2.1 (refs. 34,35). The initial model was generated in RELION. Subsequently, 664,971 good particles were further classified in 3D without symmetry. As indicated in Extended Data Fig. 3b, some classes exhibited very weak density for one or two of the six subunits. Therefore, 164,749 particles in classes with six intact subunits were re-extracted in the original pixel size of 1.49 Å/pix and refined in C3 symmetry. The overall gold-standard resolution was 4.25 Å, with the local resolution in the core TM region extending to 3.8 Å and that in the peripheral region extending to 6.5 Å (Extended Data Fig. 3c-d).

### Model building

The extracellular, transmembrane, and intracellular regions of HsLRRC8A were built *de novo* into the electron density map in COOT^36^, using the Innexin-6 (PDB ID: 5H1Q) structure as a guide. The LRR region was initially generated by creating a homology model based on the LRR protein from *L. interrogans* (PDB ID: 4U08), using the Phyre2 server ^24^. The initial model was then refined, using PHENIX^37^ with secondary structure restraints. For cross-validation, the final model was refined against one of the half maps generated in RELION, and FSC curves were calculated between the refined model and the half map 1, and between the refined model and the half map 2 (Extended Data Fig. 3e).

### Liposome reconstitution

The purified HsLRRC8A protein was incorporated into artificial liposomes, using a modified sucrose procedure^38^. Briefly, a 200 µl aliquot of a 25 mg/ml solution of azolectin in chloroform (Avanti Polar Lipids) was dried in a glass test tube under a stream of N_2_ while rotating the tube to form a homogeneous lipid film. After the lipid was dried, 5 µl of pure water was added to the bottom (pre-hydration), followed by 1 ml of 0.4 M sucrose. The solution was incubated for 3 h at 50 °C until the lipid was resuspended. After cooling the solution to room temperature, the purified HsLRRC8A protein was added to achieve the desired protein-to-lipid ratio (protein: lipid = 1:1000, w/w). The glass tube containing the protein-lipid solution was shaken gently on an orbital mixer for 3 h at 4 °C. After this procedure, the sample was ready for patch clamping.

### Patch clamping recordings

An aliquot of liposomes (2-4 µl) was taken from the liposome cloud and introduced to the recording bath. All recordings were performed with the excised liposome patched. For recording under asymmetric conditions, the pipette buffer (70mM KCl, 1.5??mM CaCl_2_, 1??mM MgCl_2_, 10 mM dextrose, and 10??mM HEPES, pH 7.4) and the bath buffer (500??mM KCl, 1.5??mM CaCl_2_, 1??mM MgCl_2_, 10 mM dextrose, and 10??mM HEPES, pH 7.4) were used. For recording under symmetric conditions, the bath and pipette buffers were identical and consisted of a solution of 70??mM KCl, 1.5??mM CaCl_2_, 1??mM MgCl_2_, 10 mM dextrose, and 10??mM HEPES, pH 7.4. The pH of buffers was adjusted by KOH. For patch pipette preparation, the borosilicate glass pipettes (BF150-86-10, Sutter Instrument, Novato) were pulled using a Flaming/Brown micropipette puller (P-97, Sutter Instrument, Novato) and polished to a diameter corresponding to a pipette resistance in the range of 4.0-6.0 MΩ. The recordings were performed when the pipette was tightly sealed with the liposome membrane and the patch resistance increased toward ∼2 GΩ. The data were acquired at 50 kHz with a 0.5-kHz filter and a 50 Hz notch filter, using an EPC-10 amplifier (HEKA, Lambrecht, Germany). All experiments were performed at room temperature (21-24 °C). All electrophysiological data were analyzed by using the pCLAMP10 software (Axon Instruments, Foster City, CA).

### Cell line formation for measurements of VRAC activity

HEK293A cells (Invitrogen) and *LRRC8A*-knockout HEK293A cells were cultured in DMEM-high glucose (Sigma-Aldrich, Cat. #D5671), supplemented with 10% fetal bovine serum (FBS; BioWest, Cat. #S1560-500) and 100 units/ml penicillin G (Meiji Seika, Cat. #6111400D2039). All cells were cultured in a 5% CO_2_ atmosphere at 37°C and verified to be negative for mycoplasma. *LRRC8A*-knockout HEK293A cells were established using the CRISPR-Cas9 system^39^. Briefly, the targeting sequence for human LRRC8A, 5’-GCACAACATCAAGTTCGACGTGG-3’, was designed using the CRISPR design tool (http://crispr.mit.edu/), and oligo DNAs for its sense and anti-sense strands were purchased from Eurofins Genomics. The sgRNA expression construct was generated by inserting the annealed guide oligo duplex with *Bbs*I into pSpCas9(BB)-2A-Puro (PX459) V2.0, a gift from Dr. Feng Zhang (Addgene plasmid #62988). The targeting sequence of the generated constructs was verified by sequencing. Subsequently, HEK293A cells were transfected with the *LRRC8A* sgRNA expression plasmid, using Lipofectamine 2000 (Invitrogen, Cat. #11668-019). After 29 h, the cells were selected by 1 µg/ml puromycin for 44 h, followed by limiting dilution to establish clonal cell lines. The generated *LRRC8A*-knockout cell lines were validated by checking for the depletion of the LRRC8A proteins and the absence of novel truncated LRRC8A proteins by western blotting.

Expression plasmids for this study were constructed by standard molecular biology techniques, and all constructs were verified by sequencing. The human LRRC8A cDNA (CDS of NM_019594.3 with the silent mutation c.1509C>T) was cloned from a cDNA pool derived from HEK293A cells and subcloned into pcDNA3. LRRC8A mutants (D102A, R103A, H104A, H105A, and Y106A) were constructed from LRRC8A wild-type (WT) and subcloned into pcDNA3 (Invitrogen). A halide-sensitive YFP (H148Q/I152L)^26^ mutant was generated from YFP (WT) and subcloned into pcDNA3.

### Western blotting and antibodies

Cells were lysed in lysis buffer (20 mM Tris-HCl, pH 7.5, 150 mM NaCl, 10 mM EDTA, 1% sodium deoxycholate, and 1% Triton X-100), supplemented with 1 mM phenylmethylsulfonyl fluoride (PMSF) and 5 μg/ml leupeptin. The cell extracts were clarified by centrifugation, and aliquots of the supernatants were mixed with 2x SDS sample buffer (80 mM Tris-HCl, pH 8.8, 80 μg/ml bromophenol blue, 28.8% glycerol, 4% SDS, and 10 mM dithiothreitol). After boiling at 98°C for 3 min, the samples were resolved by SDS-PAGE and electroblotted onto an Immobilon-P membrane (Millipore, Cat. #IPVH00010). The membrane was blocked with 2.5% skim milk in TBS-T (50 mM Tris-HCl, pH 8.0, 137 mM NaCl, and 0.05% Tween 20) and probed with the appropriate antibodies. Antibody-antigen complexes were detected on X-ray films (FUJIFILM, Cat. #47410-07523, Cat. #47410-26615 or Cat. #47410-07595) using the ECL system (GE Healthcare). Antibodies against LRRC8A (Cat. #A304-175A, 1:5,000), YFP-tag (YFP; Clone #1E4, Cat. #M048-3, 1:10,000), and actin (Cat. #A3853, 1:5,000) were purchased from Bethyl Laboratories, MBL, and Sigma-Aldrich, respectively. HRP-conjugated secondary antibodies against rabbit IgG (Cat. #7074, 1:2,000) and mouse IgG (Cat. #7076, 1:10,000–1:20,000) were purchased from Cell Signaling Technology.

### Cell-based measurements of VRAC activity

The VRAC activity in LRRC8A-rescued *LRRC8A*-knockout HEK293A cells was evaluated by the efficiency of iodide influx in the halide-sensitive YFP quenching assay^9,10,26^. *LRRC8A*-knockout HEK293A cells were transfected with YFP (H148A/I152L) and/or LRRC8A mutant expression plasmids, using Lipofectamine 2000. After 4 h, the cells were reseeded in 96-well black imaging plates (Falcon, Cat. #353219) and cultured for 65 h. Prior to measurements, the culture medium was exchanged with 50 μl/well of isoosmotic buffer (332 mOsm (kg H_2_O)–^1^, pH 7.4; 145 mM NaCl, 5 mM KCl, 2 mM CaCl_2_, 1 mM MgCl_2_, 10 mM glucose, 10 mM HEPES). The initial YFP fluorescence (Ex: 500 ± 6 nm, Em: 545 ± 6 nm) was measured 5 times at 10-sec intervals using a Varioskan Flash luminometer (Thermo Fisher Scientific). After the addition of 125 μl/well of iodide-containing hypoosmotic buffer (190 mOsm, pH 7.4; 70 mM NaI, 5 mM NaCl, 5 mM KCl, 2 mM CaCl_2_, 1 mM MgCl_2_, 10 mM glucose, 10 mM HEPES) or isoosmotic buffer (332 mOsm, pH 7.4; 70 mM NaI, 5 mM NaCl, 5 mM KCl, 2 mM CaCl_2_, 1 mM MgCl_2_, 10 mM glucose, 10 mM HEPES, 140 mM mannitol), subsequent measurements were performed 45 times at 10-sec intervals. For the data analysis, each fluorescent value was subjected to background subtraction by the fluorescent values of non-YFP (H148Q/I152L)-transfected cells, followed by normalization with the fluorescent value at the 1st time point after the addition of the iodide-containing buffer. To evaluate the efficiency of iodide influx, the normalized fluorescence values obtained after adding the iodide-containing buffer from sextuplicate wells were fitted to a one-phase exponential decay curve *y* = *A exp*(*-t/t*) + *B* by nonlinear least squares on R, using RStudio. Using the estimated parameters, the speed of YFP quenching at time *t* was calculated as *-dy/dt* = *A/t exp*(*-t/t*), and the maximum speed of YFP quenching was defined as the speed of YFP quenching at the 1st time point after the addition of the iodide-containing buffer: *A/t*. Finally, the mean of the maximum speed of YFP quenching was computed from 4 independent experiments, for the comparisons between the samples.

### Immunocytochemistry

*LRRC8A*-knockout HEK293A cells were transfected with the LRRC8A mutant expression plasmids, using Lipofectamine 2000. After 5 h, the cells were reseeded on a 1% gelatin-coated cover slip (Matsunami Glass, Cat. #C015001) and cultured for 65 h. The cells were fixed by 4% formaldehyde in PBS (137 mM NaCl, 2.7 mM KCl, 8.1 mM Na_2_HPO_4_•12H_2_O, 1.47 mM KH_2_PO_4_) for 15 min at room temperature, permeabilized with 1% Triton X-100 in PBS for 15 min at room temperature, blocked with 5% skim milk in TBS-T for 30 min at room temperature, and incubated with the primary antibody for LRRC8A overnight at 4°C. The samples were incubated with an Alexa Fluor 594-conjugated secondary antibody against rabbit IgG (Molecular Probes, Cat. #R37117) for 2 h at room temperature, followed by nuclear staining with Hoechst 33258 (Dojindo, Cat. #343-07961) for 30 min at room temperature. The cover slip was mounted on a microscope slide (Matsunami Glass, Cat. #S2444) with Fluoromount (Diagnostic BioSystems, Cat. #K024). The images were collected with a confocal microscope, TCS-SP5 (Leica), with an HCX PL APO 63x/1.40–0.60 oil objective lens.

### Statistical analysis

The data are summarized as the mean ± s.e.m. No statistical method was utilized to predetermine sample size. Homoscedasticity was assumed unless *P* < 0.01 in Levene’s test based on the absolute deviations from the median. Statistical tests and sample size *n* are indicated in the figure legends. Statistical tests were performed using R with RStudio (http://www.rstudio.com/), and *P* < 0.05 was considered statistically significant. No samples were excluded from statistical tests. The investigators were not blinded to allocation during experiments and outcome assessment. The experiments were not randomized.

### Data availability

The raw image of HsLRRC8A after motion correction has been deposited in the Electron Microscopy Public Image Archive (EMPIAR), under the accession number XXXX. The cryo-EM density map of HsLRRC8 has been deposited in the Electron Microscopy Data Bank (EMDB), under the accession number YYYY. The molecular coordinates of HsLRRC8A have been deposited in the RCSB Protein Data Bank (PDB), under the accession number ZZZZ.

## Acknowledgements

We thank the members of the Nureki lab, the Kikkawa lab, and the RIKEN Center for Life Science Technologies, especially Dr. Mizuki Takemoto and Dr. Reiya Taniguchi (Nureki lab) for assistance with the manuscript preparation and Dr. Hideki Shigematsu (RIKEN) for microscope maintenance. We also thank Dr. Yuhei Araiso (Kyoto Sangyo University) for instructions regarding the digitonin preparation; Mr. Kouki Touhara (The Rockefeller University) for advice about the anti-GFP nanobody preparation; and Dr. Kenji Iwasaki, Dr. Naoyuki Miyazaki, and Dr. Akihiro Kawamoto (Osaka University) for optimization of the cryo-EM experiment. This work was supported by a grant from the National Key R&D Program of China (2016YFA0502800); by the RIKEN Pioneering Project “Dynamic Structural Biology”; by the Platform for Drug Discovery, Informatics and Structural Life Science from Japan Agency for Medical Research and Development (AMED) to O.N.; by AMED (Grant No. JP17gm5010001) to H.I.; by a MEXT Grant-in-Aid for Specially Promoted Research (Grant No. 16H06294) to O.N.; by the National Natural Science Foundation of China (Grant No. 31571083) to Z.Y.; by the Program for Professor of Special Appointment (Eastern Scholar of Shanghai, TP2014008) to Z.Y.; by the Shanghai Rising-Star Program (4QA1400800) to Z.Y.; by the Young 1000 Talent Program of China to Z.Y.; and by JSPS KAKENHI (Grant Nos. 17J06101 to G.K.; 17K15086 to K.W.; 25221302 to H.I.; 17H03640 to R.I.; 24227004 to O.N.). This research was also supported by the Platform Project for Supporting Drug Discovery and Life Science Research (Basis for Supporting Innovative Drug Discovery and Life Science Research (BINDS)) from AMED.

## Author contributions

G.K. and O.N. conceived and designed the project. G.K. screened and purified the protein, and prepared cryo-EM samples. N.D. and R.N. assisted with the preparation of cryo-EM samples. G.K., T.Nakane, T.Y., and M.S. performed cryo-EM data collection and processing. T.Nishizawa, T.K., N.M., A.T., H.Y., and M.K. assisted with the optimization of the cryo-EM data collection. G.K. and R.I. performed the model building, with assistance from T.Nishizawa. Y.J., M.H., and Z.Y. performed the patch clamping recordings. M.I., K.W., and H.I. performed the cell-based quenching assay. G.K., R.I., and O.N. wrote the manuscript. O.N. supervised all of the research.

## Extended Data Figure Legends

**Extended Data Figure 1.**
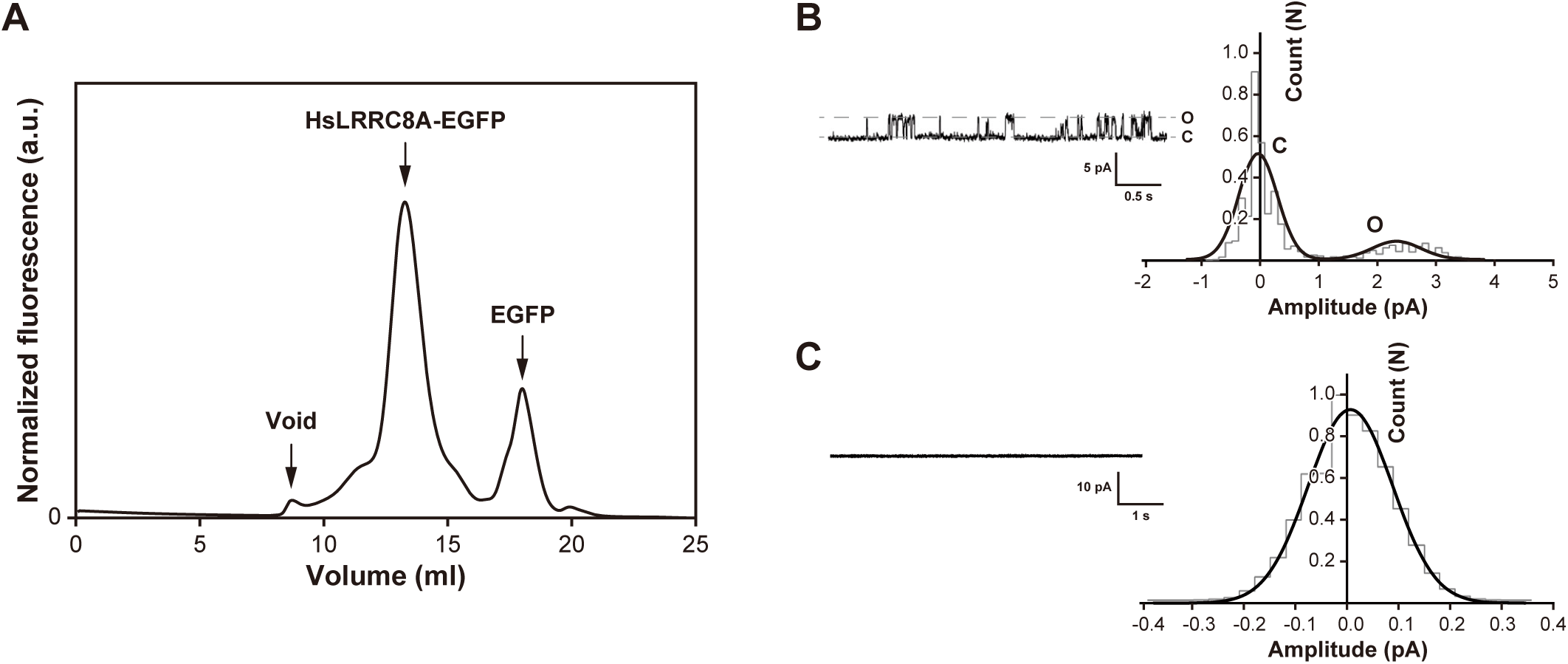
Functional properties of the HsLRRC8A protein. (**a**) FSEC profiles on a Superose 6 Increase 10/300 GL column (GE Healthcare) for the EGFP-fused HsLRRC8A expressed in HEK293S GnTI^−^ cells. The arrows indicate the estimated elution positions of the void volume, the EGFP-fused HsLRRC8A, and the free EGFP. (**b**) Single-channel current recordings of HsLRRC8A after reconstitution in liposomes under asymmetric salt conditions (500 mM KCl in bath, 70 mM KCl in pipette), and the normalized all-points amplitude histogram analysis of HsLRRC8A in the open and closed states at 100 mV. The gray dashed lines indicate the closed (C) and open (O) states. For the plot, the bin width is set at 0.1 pA/bin and the total count of events is normalized to 1.0. The distribution data were fit by the sum of two Gaussians, and the peaks correspond to the closed (C) and open (O) states. (**c**) Single-channel current recordings of HsLRRC8A after reconstitution in liposomes under symmetric salt conditions (70 mM KCl in bath, 70 mM KCl in pipette), and the normalized all-points amplitude histogram analysis of HsLRRC8A at 100 mV.

**Extended Data Figure 2.**
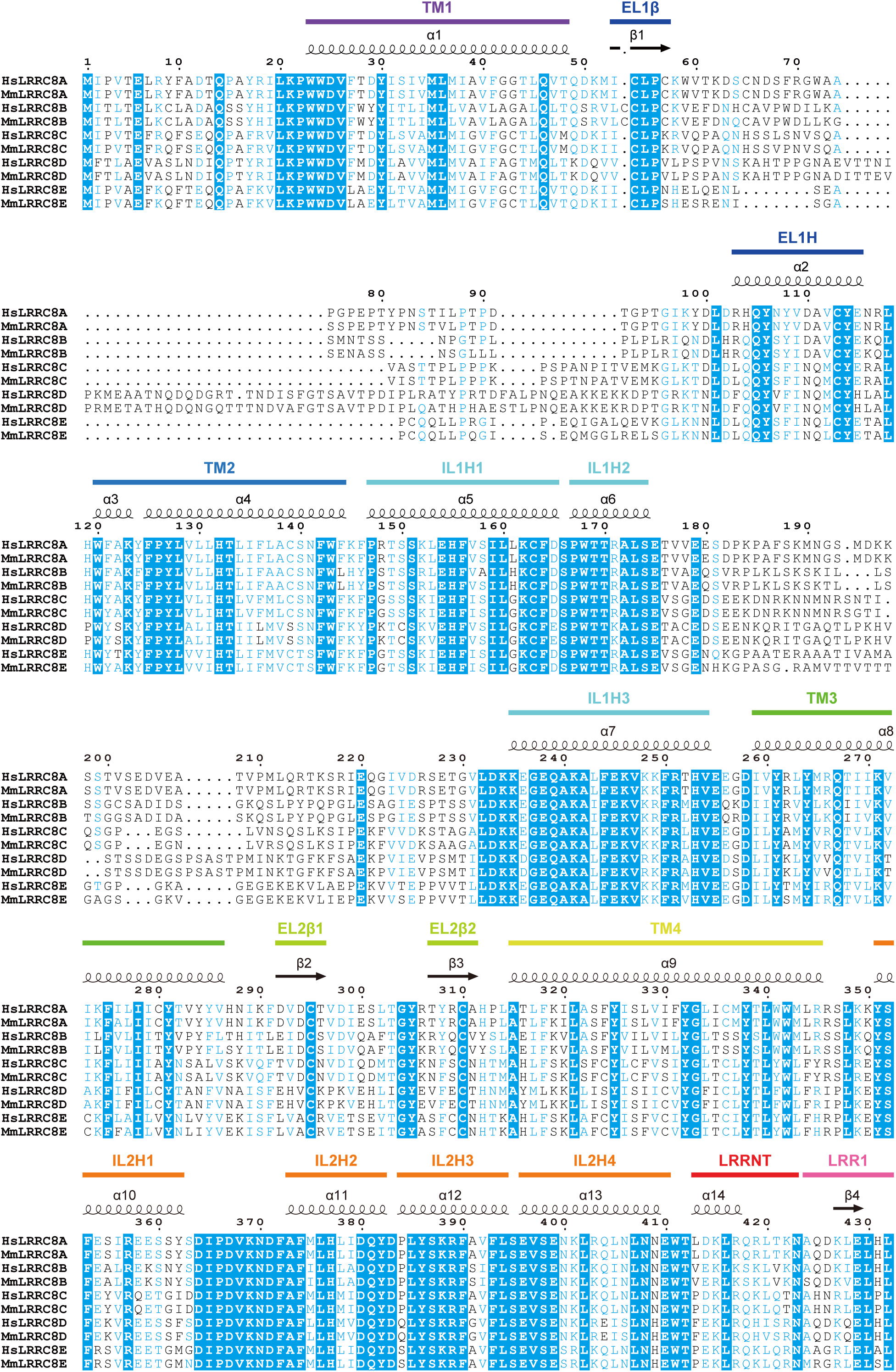
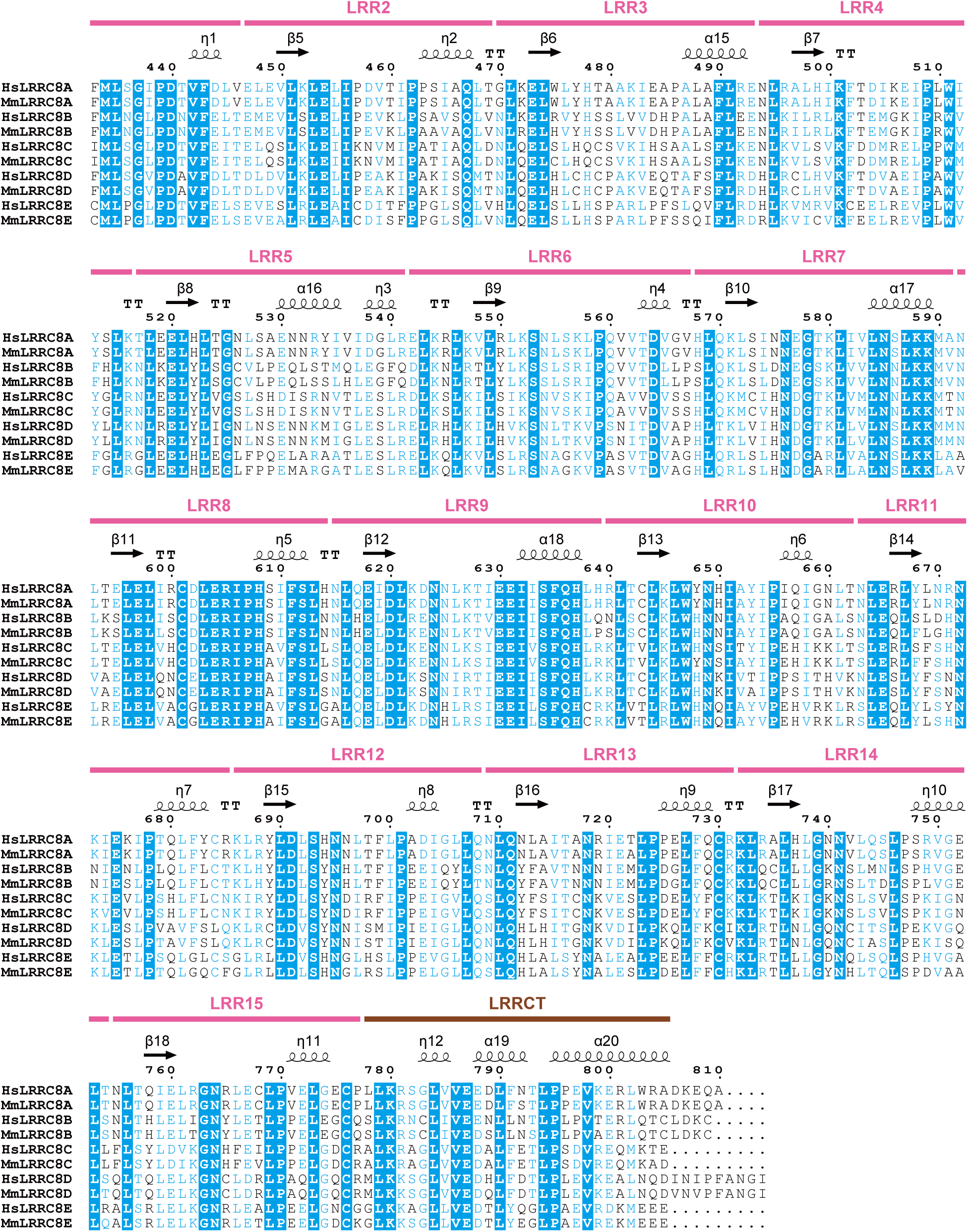
Sequence alignment of LRRC8 isoforms. The amino acid sequences of LRRC8 isoforms were aligned using Clustal Omega^41^, and are shown using ESPript3 (ref. 42). The secondary structure elements and the colors of the domains from HsLRRC8A are labeled above the alignments. For the sequence alignment, the human and mouse LRRC8 isoforms were used: human (HsLRRC8A, NCBI Reference sequence number: NP_062540.2; HsLRRC8B, NP_056165.1; HsLRRC8C, NP_115646.2; HsLRRC8D, NP_060573.2; HsLRRC8E, AAH70089.1) and mouse (MmLRRC8A, NP_808393.1; MmLRRC8B, NP_001028722.1; MmLRRC8C, NP_598658.1; MmLRRC8D, NP_848816.3; MmLRRC8E, NP_082451.2).

**Extended Data Figure 3.**
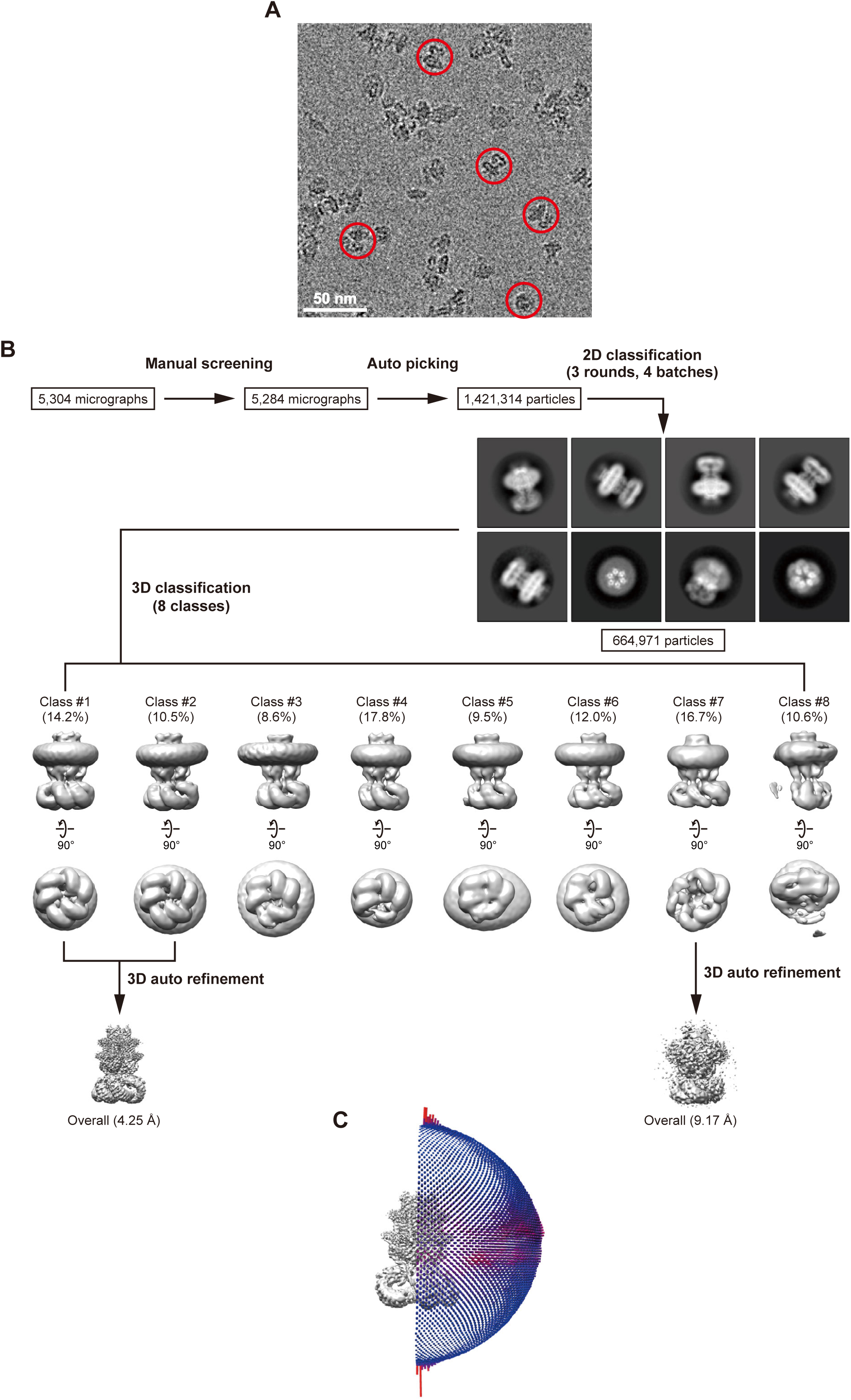
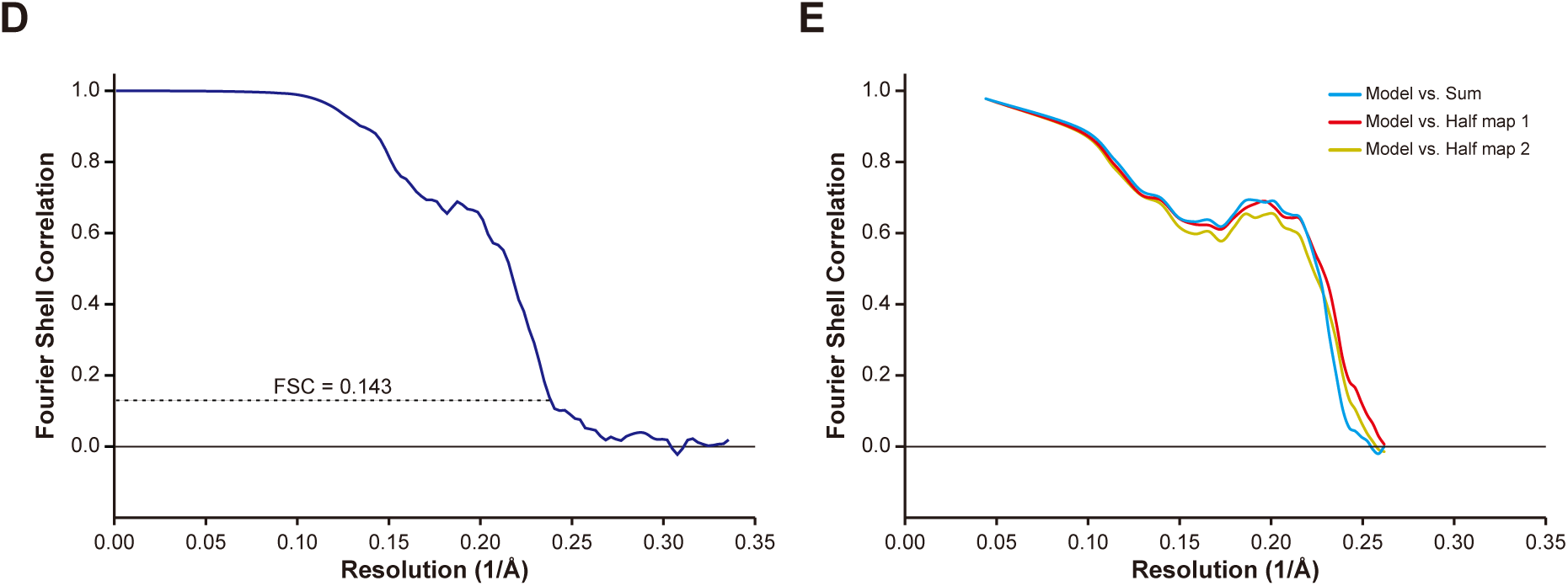
Cyro-EM analysis of HsLRRC8A. (**a**) A representative cryo-EM micrograph of HsLRRC8A. Red circles indicate individual particles. The white scale bar represents 50 nm. (**b**) Flow chart of cryo-EM data processing of the HsLRRC8A structure, including particle picking, classification, and 3D refinement. (**c**) Angular distribution plot of particles included in the final 3D reconstruction of HsLRRC8A, with C3 symmetry imposed. (**d**) Fourier Shell Correlation (FSC) curve of the final 3D reconstruction model calculated using “relion_postprocess” with masked marked 4.25 Å resolution, corresponding to the FSC = 0.143 gold standard cut-off criterion. (**e**) Cross-validation FSC curves for the refined model versus half maps (Half maps 1 and 2), and versus summed map.

**Extended Data Figure 4.**
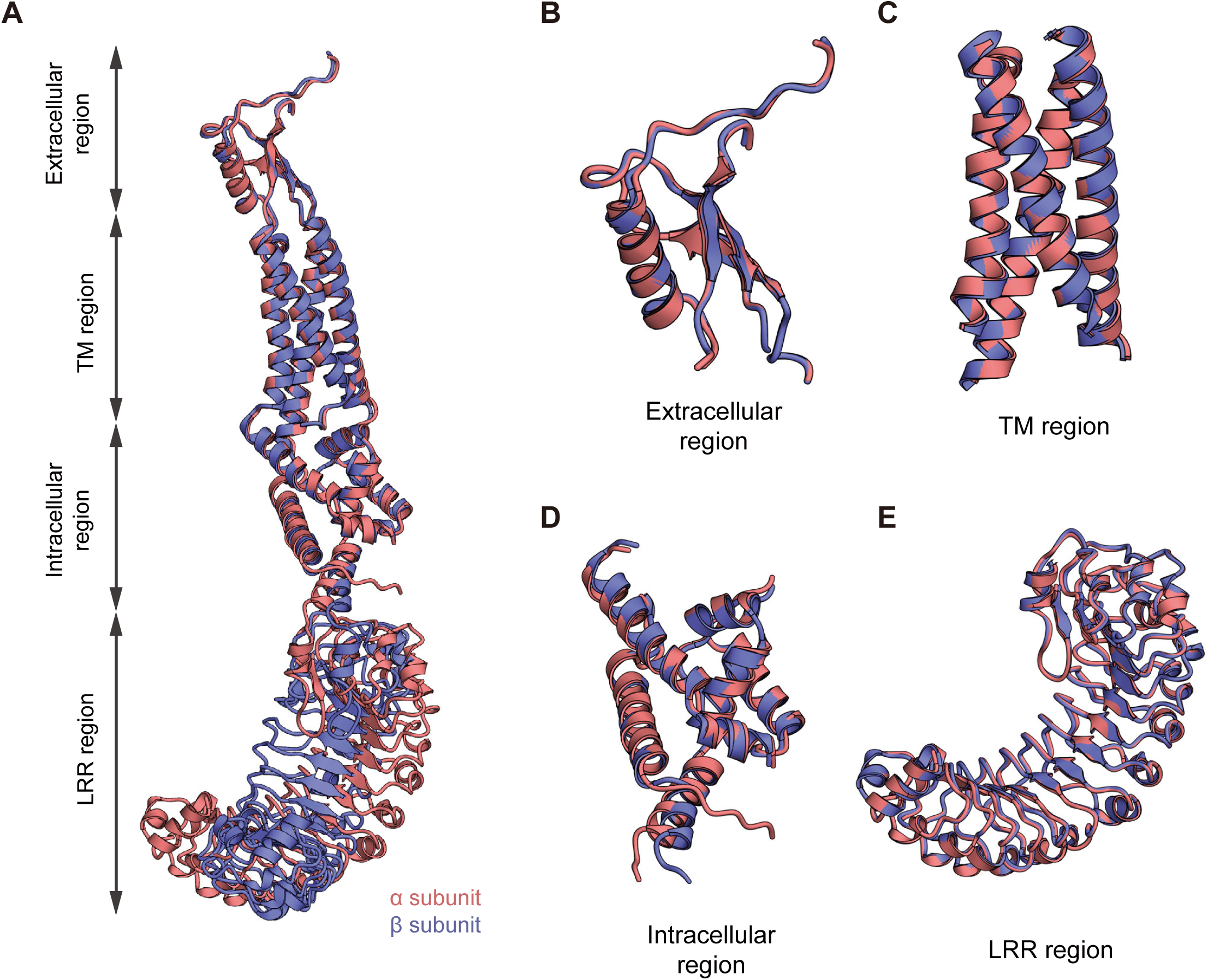
Structural comparison of two adjacent subunits. (**a**) Two adjacent subunits (α and β subunits) of HsLRRC8A are superimposed on each other. The RMSD value for the 553 Cα atoms from the subunits is 3.79 Å. For clarity, the overall subunits superimposed using only the TM region are presented. (**b**) The extracellular regions from two adjacent subunits are superimposed on each other. The RMSD value for the 68 Cα atoms from the two extracellular regions is 0.39 Å. (**c**) The transmembrane regions from two adjacent subunits are superimposed on each other. The RMSD value for the 110 Cα atoms from the two extracellular regions is 0.40 Å. (**d**) The intracellular regions from two adjacent subunits are superimposed on each other. The RMSD value for the 119 Cα atoms from the two extracellular regions is 0.98 Å. (**e**) The LRR regions from two adjacent subunits are superimposed on each other. The RMSD value for the 395 Cα atoms from the two extracellular regions is 0.47 Å.

**Extended Data Figure 5.**
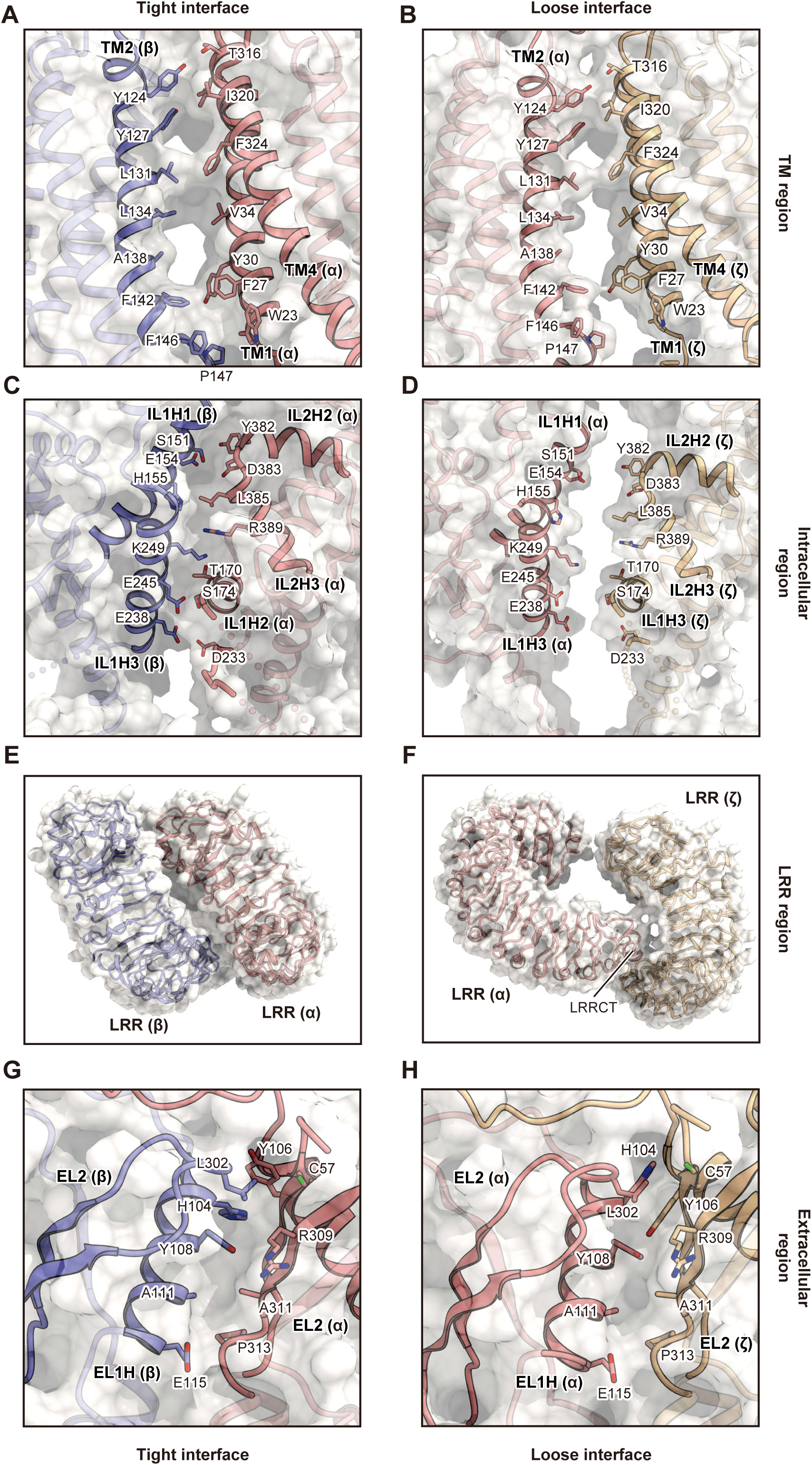
Subunit interfaces between two adjacent subunits. (**a,b**) Close-up views between two adjacent subunits at the extracellular region, showing the tight (a) and loose (b) interactions. (**c,d**) Close-up views between two adjacent subunits at the transmembrane region, showing the tight (c) and loose (d) interactions. (**e,f**) Close-up views between two adjacent subunits at the intracellular region, showing the tight (e) and loose (f) interactions. (**g,h**) Close-up views between two adjacent subunits at the LRR region, showing the tight (g) and loose (h) interactions. A surface model with the side chains of the residues lining the subunit interfaces depicted by stick models is presented for each panel. Three adjacent subunits (α, β, and ζ subunits) of HsLRRC8A are colored according to Fig. 1d.

**Extended Data Figure 6.**
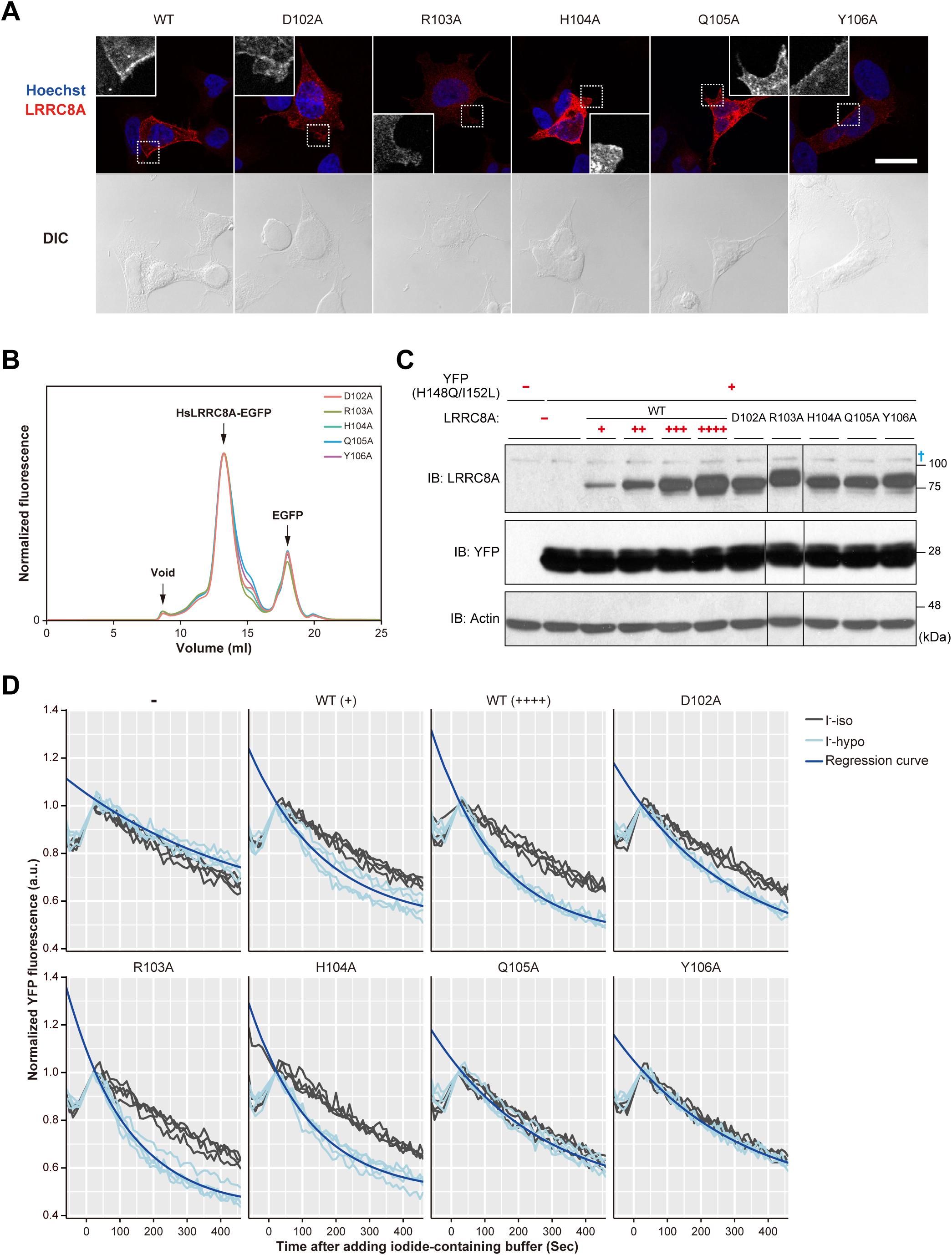
Validation of LRRC8A mutants and YFP quenching assay. (**a**) Subcellular localization of LRRC8A mutants rescued in *LRRC8A*-knockout HEK293A cells. The white dashed square indicates a region around the cell membrane; the magnified image of the LRRC8A-derived fluorescent channel is located in the corner. The white scale bar indicates 25 μm. (**b**) FSEC profiles on a Superose 6 Increase 10/300 GL column (GE Healthcare) for the EGFP-fused HsLRRC8A mutants expressed in HEK293S GnTI^−^ cells. The arrows indicate the estimated elution positions of the void volume, the EGFP-fused HsLRRC8A, and the free EGFP. The analysis of the mutants showed symmetric and monodisperse peaks, similar to those of the wild-type channel, confirming the proper structural integrities of these mutant channels. (**c**) Amounts of HsLRRC8A expression in rescued *LRRC8A*-knockout HEK293A cells. † indicates non-specific bands. (**d**) Comparison between individual time courses and the fitting curve of the normalized YFP fluorescence. Grey lines and light blue lines indicate individual time courses of normalized YFP fluorescence (*n* = 4). Blue lines indicate the mean of one-phase exponential decay-fitting curves. I^−^-iso: 332 mOsm, I^−^-hypo: 231 mOsm.

**Extended Data Figure 7.**
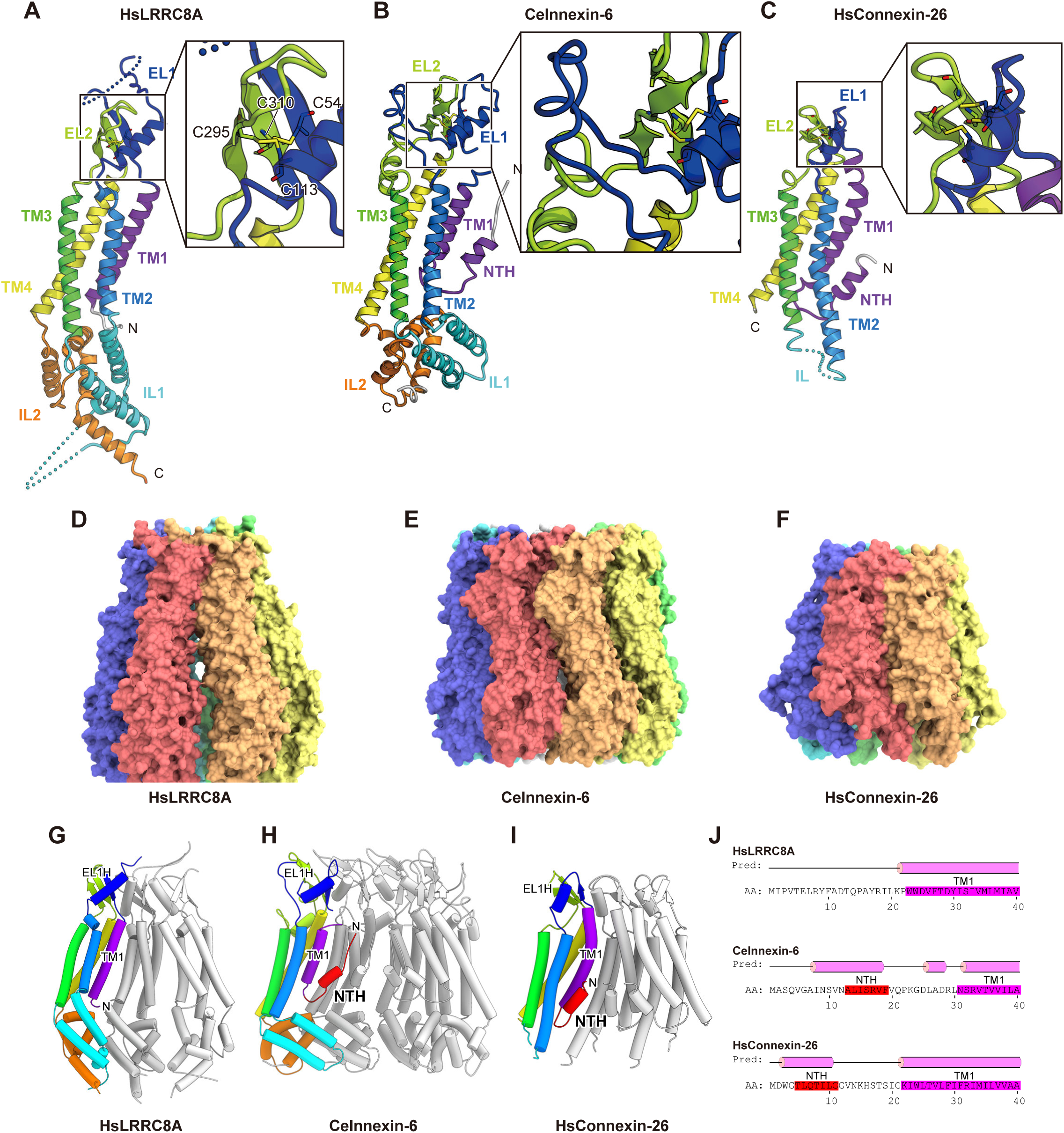
Structural similarity between HsLRRC8A, innexin, and connexin. (**a-c**) The N-terminal halves of the HsLRRC8A subunit (a), the CeInnexin-6 (PDB ID: 5H1Q) subunit (b), and the HsConnexin-26 (PDB ID: 2ZW3) subunit (c) are presented. The subunits of CeInnexin-6 and HsConnexin-26 are colored according to the HsLRRC8A coloring. The N-and C-termini are indicated by ‘N’ and ‘C’, respectively. In each structure, the Cys residues forming disulfide bonds at the extracellular region are depicted by stick models. In each panel, a close-up view of the disulfide bonds is shown in the inset. (**d-f**) The solvent-excluded molecular surfaces of HsLRRC8A (d), CeInnexin-6 (e), and HsConnexin-26 (f). The surfaces are colored according to the subunits. For HsLRRC8A, the extracellular, TM, and intracellular regions are shown. (**g-i**) The channel pore and N-terminal region of HsLRRC8A (d), CeInnexin-6 (e), and HsConnexin-26 (f). The protein main chains are shown as cylinder models. (**j**) The secondary structure prediction of the N-terminal regions of HsLRRC8A, CeInnexin-6, and HsConnexin-26. The prediction was performed by the program PSIPRED^43^. The actual locations of the corresponding helices are indicated by the red and pink boxes for NTH and TM1, respectively.

**Extended Data Figure 8.**
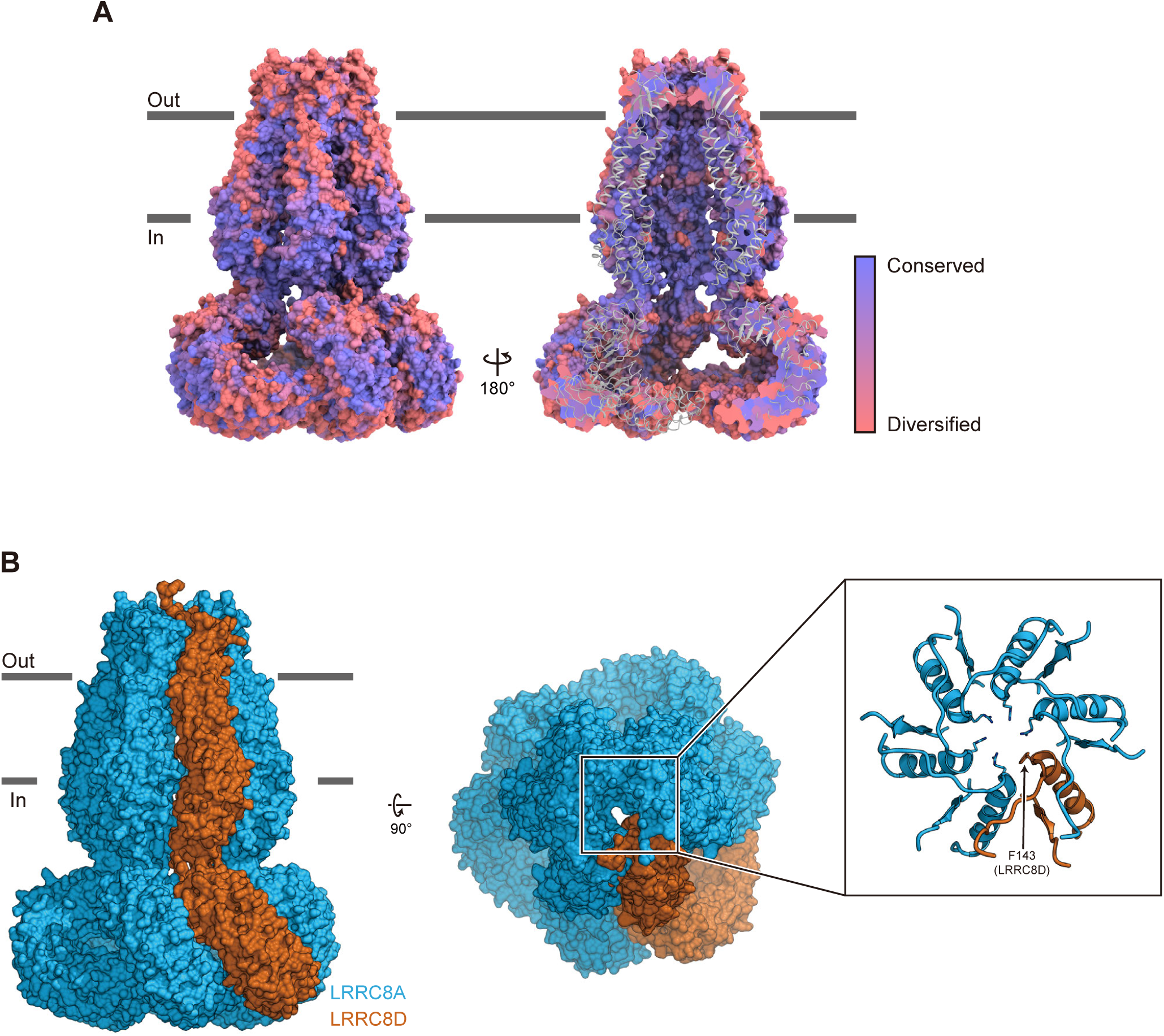
Model of the heteromeric LRRC8 protein. (**a**) The conservation score mapped on the molecular surface of HsLRRC8A. The score was calculated by the program CONSURF^27^, based on the multiple-sequence alignment of the A-E isomers in Extended Data Fig. 2. The molecular surface is colored according to the normalized conservation score, from 0.752 (pink) to −0.983 (purple). (**b**) The model structure of the (LRRC8A)_5_·LRRC8D heterohexamer, viewed parallel to the membrane (left), and from the extracellular side (right). The homology model of human LRRC8D was generated by the program MODELLER^44^, using the alignment in Extended Data Fig. 2, and then the α subunit of the present structure was replaced by this model. In the inset, a close-up view of the channel pore forming regions from the extracellular side is shown, and the side chains of pore-lining amino residues are depicted in stick representations. The five LRRC8A subunits are colored light blue, while the LRRC8D isoform is colored light brown.

**Extended Data Table 1.** Data collection, refinement and validation statistics.

|  |  |
| --- | --- |
| **Data collection** |  |
| Microscope | FEI Tecnai Arctica |
| Voltage (kV) | 200 |
| Detector | Gatan K2 Summit |
| Pixel size (Å)* | 1.490 |
| Nominal magnification* | 23,500x |
| Electron dose (e<sup>-</sup>/Å<sup>2</sup>)* | 50 |
| Defocus range (mm) | -0.5 ~ -3.5 |
| Number of collected micrographs | 5304 |
| Number of selected micrographs | 5284 |
| **Reconstitution** |  |
| Number of used particles | 164,749 |
| Symmetry imposed | C3 |
| Resolution | 4.25 |
| **Refinement** |  |
| Number of protein residues | 4,227 |
| R.m.s. deviations |  |
| Bond lengths (Å) | 0.006 |
| Bond angles (°) | 1.124 |
| Ramachandran (%) |  |
| Favored | 88.74 |
| Allowed | 11.26 |
| Outlier | 0.00 |

\* Calibrated pixel size at the detector

## References

1. Hoffmann, E. K., Lambert, I. H. & Pedersen, S. F. Physiology of cell volume regulation in vertebrates. Physiol. Rev. 89, 193–277 (2009).

2. Pedersen, S. F., Kapus, A. & Hoffmann, E. K. Osmosensory Mechanisms in Cellular and Systemic Volume Regulation. J. Am. Soc. Nephrol. 22, 1587–1597 (2011).

3. Hoffmann, E. K. et al. Role of volume-regulated and calcium-activated anion channels in cell volume homeostasis, cancer and drug resistance. Channels 6950, 380–396 (2015).

4. Okada, Y. et al. Receptor-mediated control of regulatory volume decrease (RVD) and apoptotic volume decrease (AVD). J. Physiol. 532, 3–16 (2001).

5. Jentsch, T. J. VRACs and other ion channels and transporters in the regulation of cell volume and beyond. Nat. Rev. Mol. Cell Biol. 17, 293–307 (2016).

6. Stauber, T. The volume-regulated anion channel is formed by LRRC8 heteromers-molecular identification and roles in membrane transport and physiology. Biol. Chem. 396, 975–990 (2015).

7. Pedersen, S. F., Klausen, T. K. & Nilius, B. The identification of a volume-regulated anion channel: An amazing Odyssey. Acta Physiol. 213, 868–881 (2015).

8. Sawada, A. et al. A congenital mutation of the novel gene LRRC8 causes agammaglobulinemia in humans. J. Clin. Invest. 112, 1707–1713 (2003).

9. Qiu, Z. et al. SWELL1, a plasma membrane protein, is an essential component of volume-regulated anion channel. Cell 157, 447–458 (2014).

10. Voss, F. K. et al. Identification of LRRC8 Heteromers as an Essential Component of the Volume-Regulated Anion Channel VRAC. Science 344, 634–638 (2014).

11. Kubota, K. et al. LRRC8 involved in B cell development belongs to a novel family of leucine-rich repeat proteins. FEBS Lett. 564, 147–152 (2004).

12. Smits, G. & Kajava, A. V. LRRC8 extracellular domain is composed of 17 leucine-rich repeats. Mol. Immunol. 41, 561–562 (2004).

13. Abascal, F. & Zardoya, R. LRRC8 proteins share a common ancestor with pannexins, and may form hexameric channels involved in cell-cell communication. BioEssays 34, 551–560 (2012).

14. Oshima, A., Tani, K. & Fujiyoshi, Y. Atomic structure of the innexin-6 gap junction channel determined by cryo-EM. Nat. Commun. 7, 13681 (2016).

15. Maeda, S. et al. Structure of the connexin 26 gap junction channel at 3.5 A resolution. Nature 458, 597–602 (2009).

16. Bennett, B. C. et al. An electrostatic mechanism for Ca2+ -mediated regulation of gap junction channels. Nat Commun 7, 8770 (2016).

17. Syeda, R. et al. LRRC8 Proteins Form Volume-Regulated Anion Channels that Sense Ionic Strength. Cell 164, 499–511 (2016).

18. Gradogna, A., Gavazzo, P., Boccaccio, A. & Pusch, M. Subunit-dependent oxidative stress sensitivity of LRRC8 volume-regulated anion channels. J. Physiol. 595, 6719–6733 (2017).

19. Planells-Cases, R. et al. Subunit composition of VRAC channels determines substrate specificity and cellular resistance to Pt-based anti-cancer drugs. EMBO J. 34, 2993–3008 (2015).

20. Schober, A. L., Wilson, C. S. & Mongin, A. A. Molecular composition and heterogeneity of the LRRC8-containing swelling-activated osmolyte channels in primary rat astrocytes. J. Physiol. 595, 6939–6951 (2017).

21. Lutter, D., Ullrich, F., Lueck, J. C., Kempa, S. & Jentsch, T. J. Selective transport of neurotransmitters and-modulators by distinct volume-regulated LRRC8 anion channels. J. Cell Sci. 130, 1122–1133 (2017).

22. Kawate, T. & Gouaux, E. Fluorescence-Detection Size-Exclusion Chromatography for Precrystallization Screening of Integral Membrane Proteins. Structure 14, 673–681 (2006).

23. Hattori, M., Hibbs, R. E. & Gouaux, E. A fluorescence-detection size-exclusion chromatography-based thermostability assay for membrane protein precrystallization screening. Structure 20, 1293–1299 (2012).

24. Kelley, L. A., Mezulis, S., Yates, C. M., Wass, M. N. & Sternberg, M. J. E. The Phyre2 web portal for protein modeling, prediction and analysis. Nat. Protoc. 10, 845–858 (2015).

25. Ullrich, F., Reincke, S. M., Voss, F. K., Stauber, T. & Jentsch, T. J. Inactivation and anion selectivity of volume-regulated anion channels (VRACs) depend on c-terminal residues of the first extracellular loop. J. Biol. Chem. 291, 17040–17048 (2016).

26. Galietta, L. J. V, Haggie, P. M. & Verkman, A. S. Green fluorescent protein-based halide indicators with improved chloride and iodide affinities. FEBS Lett. 499, 220–224 (2001).

27. Ashkenazy, H., Erez, E., Martz, E., Pupko, T. & Ben-tal, N. ConSurf 2010: calculating evolutionary conservation in sequence and structure of proteins and nucleic acids. Nucleic Acids Res. 38, W529–W533 (2010).

28. Brohawn, S. G., Campbell, E. B. & MacKinnon, R. Physical mechanism for gating and mechanosensitivity of the human TRAAK K+ channel. Nature 516, 126–130 (2014).

29. Goehring, A. et al. Screening and large-scale expression of membrane proteins in mammalian cells for structural studies. Nat. Protoc. 9, 2574–2585 (2014).

30. Kirchhofer, A. et al. Modulation of protein properties in living cells using nanobodies. Nat. Struct. Mol. Biol. 17, 133–138 (2010).

31. Mastronarde, D. N. Automated electron microscope tomography using robust prediction of specimen movements. J. Struct. Biol. 152, 36–51 (2005).

32. Zheng, S. Q. et al. MotionCor2: anisotropic correction of beam-induced motion for improved cryo-electron microscopy. Nat. Methods 14, 331–332 (2017).

33. Rohou, A. & Grigorieff, N. CTFFIND4: Fast and accurate defocus estimation from electron micrographs. J. Struct. Biol. 192, 216–221 (2015).

34. Scheres, S. H. W. Semi-automated selection of cryo-EM particles in RELION-1.3. J. Struct. Biol. 189, 114–122 (2015).

35. Kimanius, D., Forsberg, B. O., Scheres, S. H. W. & Lindahl, E. Accelerated cryo-EM structure determination with parallelisation using GPUS in RELION-2. Elife 5, 1–21 (2016).

36. Emsley, P., Lohkamp, B., Scott, W. G. & Cowtan, K. Features and development of Coot. Acta Crystallogr. D. Biol. Crystallogr. 66, 486–501 (2010).

37. Adams, P. D. et al. PHENIX: a comprehensive Python-based system for macromolecular structure solution. Acta Crystallogr. D. Biol. Crystallogr. 66, 213–21 (2010).

38. Battle, A. R., Petrov, E., Pal, P. & Martinac, B. Rapid and improved reconstitution of bacterial mechanosensitive ion channel proteins MscS and MscL into liposomes using a modified sucrose method. FEBS Lett. 583, 407–412 (2009).

39. Ran, F. A. et al. Genome engineering using the CRISPR-Cas9 system. Nat. Protoc. 8, 2281–2308 (2013).

40. Smart, O. S., Neduvelil, J. G., Wang, X., Wallace, B. A. & Sansomt, M. S. P. HOLE??: A program for the analysis of the pore dimensions of ion channel structural models. J. Mol. Graph. 14, 354–360 (1996).

41. Sievers, F. et al. Fast, scalable generation of high-quality protein multiple sequence alignments using Clustal Omega. Mol. Syst. Biol. 7, (2011).

42. Robert, X. & Gouet, P. Deciphering key features in protein structures with the new ENDscript server. Nucleic Acids Res. 42, 320–324 (2014).

43. Buchan, D. W. A. et al. Protein annotation and modelling servers at University College London. Nucleic Acids Res. 38, W563–W568 (2010).

44. Fiser, A. & Sali, A. Modeller: Generation and Refinement of Homology-Based Protein Structure Models. Methods Enzymol. 374, 461–491 (2003).

